# Injured endocardium obtains characteristics of haemogenic endothelium during adult zebrafish heart regeneration

**DOI:** 10.1101/2024.12.18.629122

**Authors:** Jun Ying, Irene Louca, Jana Koth, Abigail Killen, Konstantinos Lekkos, Zhilian Hu, Esra Sengul, William T. Stockdale, Xiaonan Wang, Mathilda T. M. Mommersteeg

**Affiliations:** School of Public Health, Shanghai Jiao Tong University School of Medicine, Shanghai, China; Institute of Developmental and Regenerative Medicine, Old Road Campus, University of Oxford, Oxford, UK; MRC Molecular Haematology Unit, Weatherall Institute of Molecular Medicine, John Radcliffe Hospital, University of Oxford, Oxford, UK; Department of Physiology, Anatomy & Genetics, University of Oxford, Oxford, UK; Faculty of Medical Laboratory Science, College of Health Science and Technology, Shanghai Jiao Tong University School of Medicine, Shanghai, China

## Abstract

Reactivation of embryonic developmental pathways during regeneration aims to restore tissue architecture and functionality. We previously reported that following cryoinjury, a heterogeneous population of Runx1-expressing endocardial cells differentially upregulates genes associate with scarring and myofibroblast identity. Further analysis of our published RNAseq data alongside 5 publicly available datasets now identifies additional heterogeneity in the Runx1-positive injured endocardium. Here, we show that the endocardium also reactivates a dormant endocardial-to-haematopoietic transition (EHT) mechanism. Runx1-expressing endocardial cells upregulate genes associated with haemogenesis and morphologically display features of EHT. Live imaging shows cells budding off the endocardium and lineage analysis identifies overlap with leukocyte markers. Ablation of *runx1* function further shifts differentiation of the endocardium towards the EHT fate. The identification of transient *runx1*-expressing cells transitioning towards myofibroblast or haemogenic endocardium identities demonstrates the complexity of the zebrafish endocardial injury response and highlights the role of Runx1 in regulating cell fate decisions in the endocardium.

## Introduction

During regeneration, tissue repair typically occurs in a manner recapitulating the developmental processes giving rise to specific organs in the embryo so that normal tissue architecture and physiological function are restored. This re-activation of developmental pathways is thought to underlie successful regeneration by initiating processes normally dormant in the adult heart to replace lost tissue, including neovascularisation and myocardial proliferation^1,2^. This process also takes place in the endocardial cells found in the wound area that upregulate pathways required for endocardial development, including retinoic acid, Wnt, TGF-β and Notch signalling^3–5^. Besides signalling to cardiomyocytes to initiate a proliferative response^3,6^, the activated endocardium undergoes morphological changes and upregulates genes involved in fibrosis, such as smooth muscle genes and collagens, thereby contributing to scar formation^5,7–9^.

Recently, we discovered that Runx1 is upregulated in a number of cell-types responding to injury in the zebrafish heart, including the endocardium^9^. *Runt*-related transcription factor-1, *Runx1* (also known as acute myeloid leukaemia 1 protein, AML1), is a key transcription factor during development and is involved in the regulation of cell proliferation and fate decisions of multiple cell-types^10–14^. Using single cell RNA sequencing (scRNA-Seq), we showed that a subset of these injured *runx1*-positive endocardial cells upregulates a myofibroblast phenotype without losing their endocardial specific gene expression, thus taking on a dual cell-type identity^9^. Up-regulation of this myofibroblast gene programme without commitment into fully mature myofibroblasts might be key to the formation of a scar that is temporary and can be degraded during successful regeneration. Removal of functional Runx1 results in a strong reduction of this dual endocardial-myofibroblast gene profile^9^. This observation fits well with recent findings in other organs, highlighting the importance of Runx1 in cell-type transition, in particular epithelial-to-mesenchymal transition (EMT) and myofibroblast formation^15–17^ and its importance for cardiac repair following injury^18–20^.

However, the control of Runx1 on cell-type transition extends beyond EMT as Runx1 is best known for its key regulatory role in endothelial-to-haematopoietic transition (EHT) during embryonic development^12,21–23^. During early stages of development, multipotent haematopoietic stem cells (HSCs) with self-renewal capability derive from endothelial cells in the ventral wall of the dorsal aorta as the definitive blood wave^24^. These HSCs give rise to different hematopoietic lineages, including myeloid and erythroid progenitors. Runx1 marks these haemogenic endothelial cells and is required for the process of EHT^24^. While EHT was initially thought to be restricted to a specific localised set of endothelial cells only in the dorsal aorta, studies have reported another site of EHT, the developing endocardium in both mouse and zebrafish embryos^25–28^. Most recently, Bornhorst et al have reported on the contribution of endocardial cells to the haematopoietic lineage supported by their observations of a sustained population of HSPCs/Megakaryocyte-Erythroid Progenitor cells in the endocardium of zebrafish embryos^28^. Therefore, the appearance of Runx1-positive endocardial cells after injury suggests the possibility of re-activation of developmental EHT in the endocardium after injury^10^.

In this study, we report that the upregulation of Runx1 in the injured endocardium can indeed be viewed as a reactivation of developmental endocardial Runx1 expression in the embryo. Detailed re-analysis of sections and published scRNA-Seq data highlights the cellular heterogeneity in the Runx1-positive injured endocardium where, in addition to a population of cells with a dual endocardial-myofibroblast phenotype, we identified a population with gene expression characteristic for haemogenic endocardium. These Runx1-positive endocardial cells in the wound acquire the morphology of haemogenic endothelial cells. Using time-lapse live imaging, we demonstrate that Kdrl-GFP endocardial cells bud off the injured endocardium. These Kdrl-positive blood cells are present in low numbers and can only be identified after injury, especially at 3dpci when the number of Runx1-positive endocardial cells in the wound also peaks. Additionally, lineage tracing analysis of the injured endocardium using the scRNA-Seq data as well as a *Tg(fli1-creERT2);Tg(ubiCSY)* line, provides further evidence that EHT occurs. We previously published that absence of functional Runx1 results in a strong reduction in the number of re-activated endocardial cells and these cells fail to transition towards the myofibroblast phenotype. We now find that in absence of *runx1*, the remaining cells favour a transition towards the EHT gene program with appearance of a mutant specific population of Kdrl-positive blood cells. These data suggest that an EHT gene programme is indeed re-activated in the injured adult endocardium with Runx1 playing an important role in lineage decisions in the endocardium. Runx1 involvement in the regulation of developmental pathways could have important implications in understanding the regenerative processes of the heart.

## Results

### The injured endocardium is highly heterogeneous after injury

During fish heart regeneration, many developmental pathways typically upregulated in the aorta of the developing embryo are re-activated in the injured endocardium, including Notch, retinoic acid, Wnt and TGF-β signalling pathways^3–5,23^. Upregulation of the same pathways results in the activation of the transcription factor Runx1 during development, which is required for the formation of the haemogenic endothelium^2,10,12,21,23^. In our previous study^9^, we noted a temporary upregulation of Runx1 in the endocardium following cardiac injury using the *TgBAC(runx1P2:Citrine)* reporter line^29^ crossed to the *Tg(kdrl:Hsa.HRAS-mCherry)* line, which labels the endothelium and endocardium. To test if the injury-specific expression is a re-activation of aspects of embryonic development, we examined the expression of Runx1 in the embryonic heart using the same Runx1:Citrine; Kdrl-mCherry fish line. Analysis of the endocardium for Runx1:Citrine expression during development showed that while there were few Citrine/mCherry double-positive cells at 2 days post fertilisation (dpf), there was a strong increase at 5dpf with expression decreasing again by 7dpf, confirming expression of Runx1 in a scattered population of endocardial cells during early heart development (Figure 1A-B). Therefore, expression after injury can indeed be considered a re-activation of embryonic expression, which raises the possibility of EHT occurring after injury.

**Figure 1.**
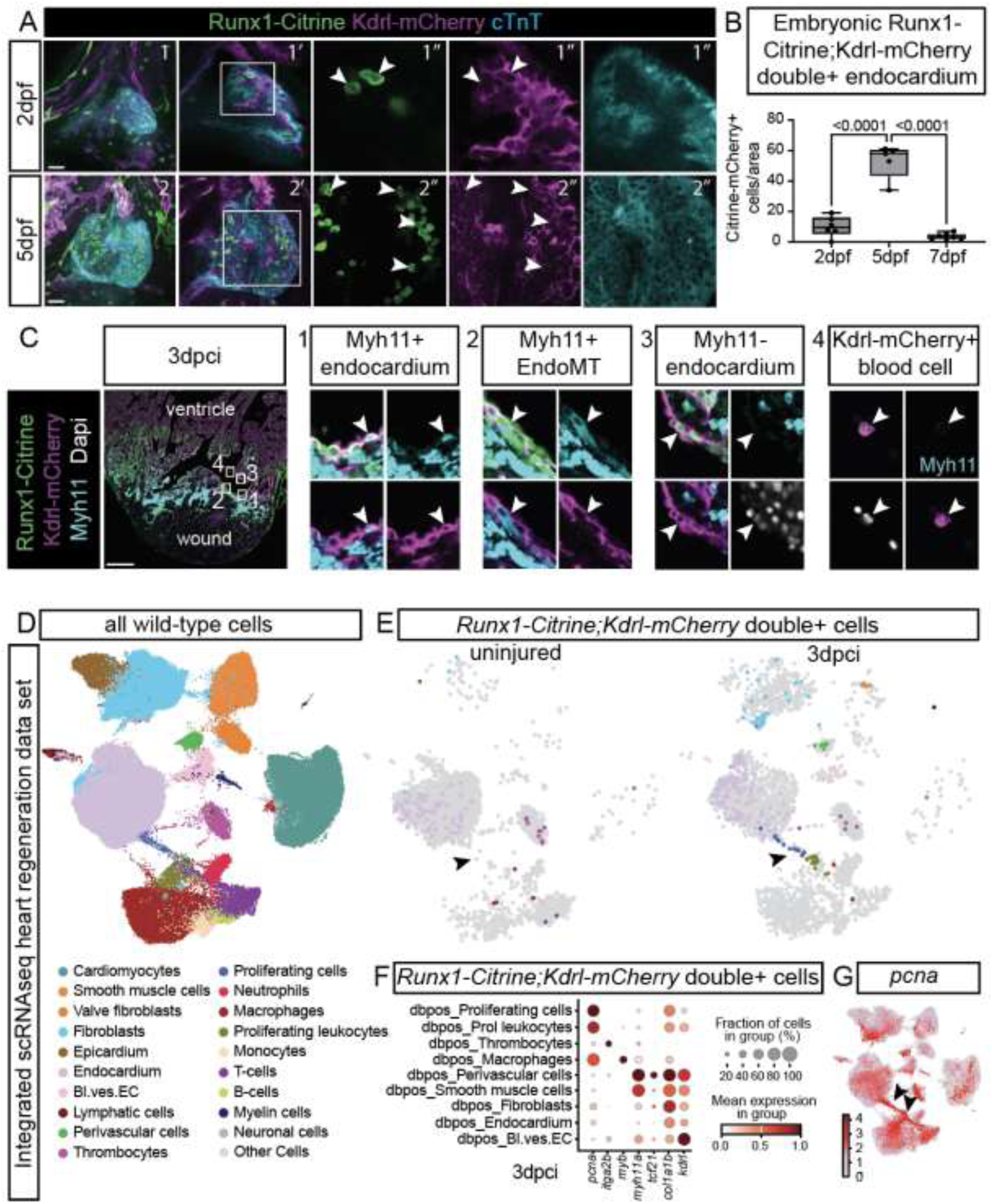
The injured endocardium is highly heterogeneous after injury. A, Immunohistochemistry of Runx1:Citrine, Kdrl-mCherry and cardiac TroponinT in the embryonic heart at 2 (images 1-1’’) and 5 dpf (images 2-2’’). Images 1-2 display maximum intensity projections in a ventral view, images 1’ -1’’, 2’-2’’ display single z-planes B, Comparison of Tg(kdrl:HsHRAS-mCherry);Tg(BAC-Runx1P2:Citrine) double positive cells in the ventricle outlined by cardiac TroponinT as shown in A. n=6 2dpf, n=5 5dpcf, n=7 7dpf, one-way ANOVA with Tukey’s test. C, Immunohistochemistry showing labelling of Runx1:Citrine, Kdrl-mCherry, Myh11 and DAPI in an adult heart at 3dpci. Numbered blocks are enlarged. Arrowheads in block 1 point to scattered Myh11-positive cells within the Runx1:Citrine positive endocardium. Arrowheads in block 2 point to Myh11-positive endocardial cells showing an EndoMT phenotype. Arrowheads in block 3 point to rounded Runx1-positive endocardial cells that are negative for Myh11. Arrowheads in block 4 point to Kdrl-mCherry positive circulating cell. D, UMAP of the integrated zebrafish heart regeneration landscape from the integration of 6 independent scRNAseq datasets^9,30–34^. E, UMAP of *runx1:citrine;kdrl-mcherry* double positive cells from the Koth et al^9^ data highlighted in the integrated landscape. Arrowheads point to the double positive cells in the proliferating cluster connecting the endocardial cluster to the leukocyte cluster after injury. F, Dotplot displaying expression of key cell identity markers in double positive *runx1-kdrl* cells in the indicated populations at 3dpci. The proliferating cluster expressed high levels of *pcna,* but also still expressed *kdrl*. The neighbouring double positive cells in the leukocyte cluster are also highly proliferative as indicated by *pcna* expression. G, UMAP of *pcna* expression shows the highly proliferative cell population connecting to the endocardial cluster and proliferating leukocyte cluster on either side. Scale bar: 100µm.

The endocardium becomes a heterogeneous cell population after injury with a subset of cells upregulating Runx1 expression^9^. Notably, within the Runx1-positive cell population, we identified further heterogeneity. As previously reported, a portion of cells upregulated a Myh11-positive myofibroblast gene programme^9^, however, only in rare instances Myh11-positive endocardial cells also displayed an endocardial to mesenchymal transition (EndoMT) phenotype with signs of invasion of the underlying cells (Figure 1C blocks 1-2). We further confirmed that there was little evidence of these cells fully undergoing EndoMT despite the upregulation of key genes for mesenchymal transformation (Supplementary Figure 1A-C). However, many of the injured Runx1-positive endocardial cells were negative for Myh11 gene expression and displayed a more rounded morphology than the Myh11-positive cells (Figure 1C block 3). Such heterogeneity as present within the same tissue section (highlighted in Figure 1C) underlines the possibility that a subset of cells could activate EHT genes, especially in light of the presence of Kdrl-mCherry-positive cells resembling circulating cells (Figure 1C block 4).

Given the assumption that EHT may be a rare event and thus requiring a large number of cells for observation, our previously published scRNAseq data set (Koth *et al*) is well-suited for analysing this possibility^9^. In our dataset, uninjured and 3dpci cells from the Runx1-Kdrl fish line were FACS sorted as Runx1:Citrine+, Kdrl-mCherry+ and Citrine-mCherry double positive in equal proportions to specifically enrich for the relatively sparse double-positive population. Furthermore, to increase the depth of the analysis and relate the double-positive cell population to all other cell types, we have integrated our data set together with 5 other publicly available scRNAseq data sets^30–34^ into one large integrated zebrafish heart regeneration data set with 279,384 cells in total, including cells from uninjured and injured fish up to 30dpci. This large high-resolution data set allowed to confidently identify 19 distinct cell-types present in the healthy or the injured heart (Figure 1D, Supplementary figure 1D). We then coloured the 357 wild-type *runx1:citrine*, *kdrl-mcherry* double positive cells onto the integrative UMAP for the uninjured and 3dpci samples, respectively. As expected, double-positive cells were mainly found in the endocardial cluster in the uninjured condition. However, after injury, subsets of double-positive cells displayed high heterogeneity and were located sparsely in a large range of cell type clusters, while retaining some level of *kdrl* expression (Figure 1E-F). One subset of double-positive cells was present in the vasculature, in both the endothelial lining as well as in *tcf21*-positive perivascular cells, which we confirmed by microscopy (Figure 1E-F, Supplementary Figure 1E). Another large sub-set of double-positive highly proliferative cells was present as a bridge on the UMAP connecting the endocardial cells to the large leukocyte cluster which only appeared after injury, peaking at 3dpci (Figure 1E, arrowheads).

Considering the possible activation of EHT following cardiac injury, this direct connection between the endocardial and leukocyte clusters via proliferating cells is interesting. The bridging cells were enriched with *pcna* expression (Figure 1G), which is a key marker gene for cell proliferation^35^. Others have described that endothelial cells need to re-enter the cell cycle for EHT to occur^36.^ If the cell cycle is blocked, haemogenic endothelial cells lose their ability to undergo this transitional process. The endocardium is known to be highly proliferative after injury^4^, and the overlap of *pcna* expression with Runx1:Citrine in the endocardium confirms this occurs within the Runx1-positive cells (Supplementary Figure 1F)^9^. The large number of double positive cells within the highly proliferative endocardium-leukocyte bridging cluster, with a subset present in the directly neighbouring leukocyte population is highly suggestive of a transitionary state between the endocardium and leukocyte populations. Therefore, Runx1-positive endocardial cells can upregulate gene programmes involved in either myofibroblast or perivascular cell differentiation, but also a transcriptional programme suggestive of the possible transition towards a leukocyte phenotype.

### The injured endocardium gains features of endothelial-to-haematopoietic transition upon injury

A number of recent scRNAseq studies on the process of EHT have identified that the cell cycle status discriminates the haemogenic endothelium from its progenitors. As cells transit from an endothelial state towards a haematopoietic state, they lose expression of endothelial genes and enter the cell cycle^37–40^. Therefore, we hypothesised that the highly proliferative cell cluster connecting the endocardial population with the also highly proliferative leukocytes could represent EHT. While we observed three additional UMAP bridges between the endocardial and blood cell clusters, these did not contain double positive *Runx1:Citrine;Kdrl-mCherry* cells and detailed analysis indicated they did not warrant further attention in this study (Figure 1D-E, Supplementary Figure 2). To gain a better understanding of the highly proliferative cells in the bridge of interest and potential haematopoietic stem and progenitor cells (HSPCs), we isolated the bridging cells and performed further sub-clustering analysis (Figure 2A-B) to identify the key cellular processes and signalling pathways, characterising each sub-cluster (Figure 2C). Clusters 1 and 2 that were located within or close to the large endocardial cluster were represented by processes including *vasculature development* and *blood vessel morphogenesis* and were enriched for genes involved in *hematopoietic progenitor cell differentiation*, while still expressing endocardial genes such as *kdrl, fli1a* and *cdh5* (Figure 2C-D). These gene expression profiles indicate the similarity of these clusters to the pre-haemogenic endothelium as seen during development^37^. Clusters 3 and 4 were largely located within the highly *pcna*-positive UMAP bridge and connecting leukocyte cluster (Figure 2D). The proliferative status of clusters 3 and 4 was reflected by processes enriched for *cell cycle,* which is characteristic of the haemogenic endothelium itself as well as HSPC^37^. Mapping specific genes of interest onto the bridging cells in question shows that *kdrl* levels decreased from cluster 1 to 4, while *pcna* levels increased (Figure 2D). Clusters 1 and 2 expressed genes such as *gata2a* and *tal1*, which are key initiators of EHT. Cluster 4, which was enriched for genes involved in processes for *“RUNX1 regulates transcription of genes involved in differentiation of HSCs”*, had low levels of expression of endocardial genes, but high levels of genes like *myb, gata2b* and *spi1b*, suggesting this cluster has HSPC-like characteristics (Figure 2C-D)^37^*. myb* and *gata2b* are also required for myeloid and lymphocyte differentiation ^41,42^, and expression of *sla2* in the HSPC-like cluster suggests differentiation towards this lineage^43^. Some cells expressed the macrophage marker *mpeg1.1*, while others expressed the neutrophil marker *mmp9,* suggesting differentiation towards several different blood lineages in this highly proliferative cluster. While *tal1* was expressed in the cells resembling pre-haemogenic endothelium, we did not note any expression in the cells possibly undergoing EHT, suggesting no evidence of erythrocyte differentiation of these cells. To investigate the scRNAseq findings, we analysed the 3dpci Runx1-positive endocardium for expression of transcription factors which are known to be key to the process of EHT*, gata2b, myb* and *tal1* (Figure 2E). We could indeed detect these genes in the Runx1:Citrine positive endocardium. These data fit well with the knowledge that haemogenic endothelium gives rise to multi-lineage haematopoietic populations^44^ and provides further evidence that haemogenesis in the endocardium could occur.

**Figure 2.**
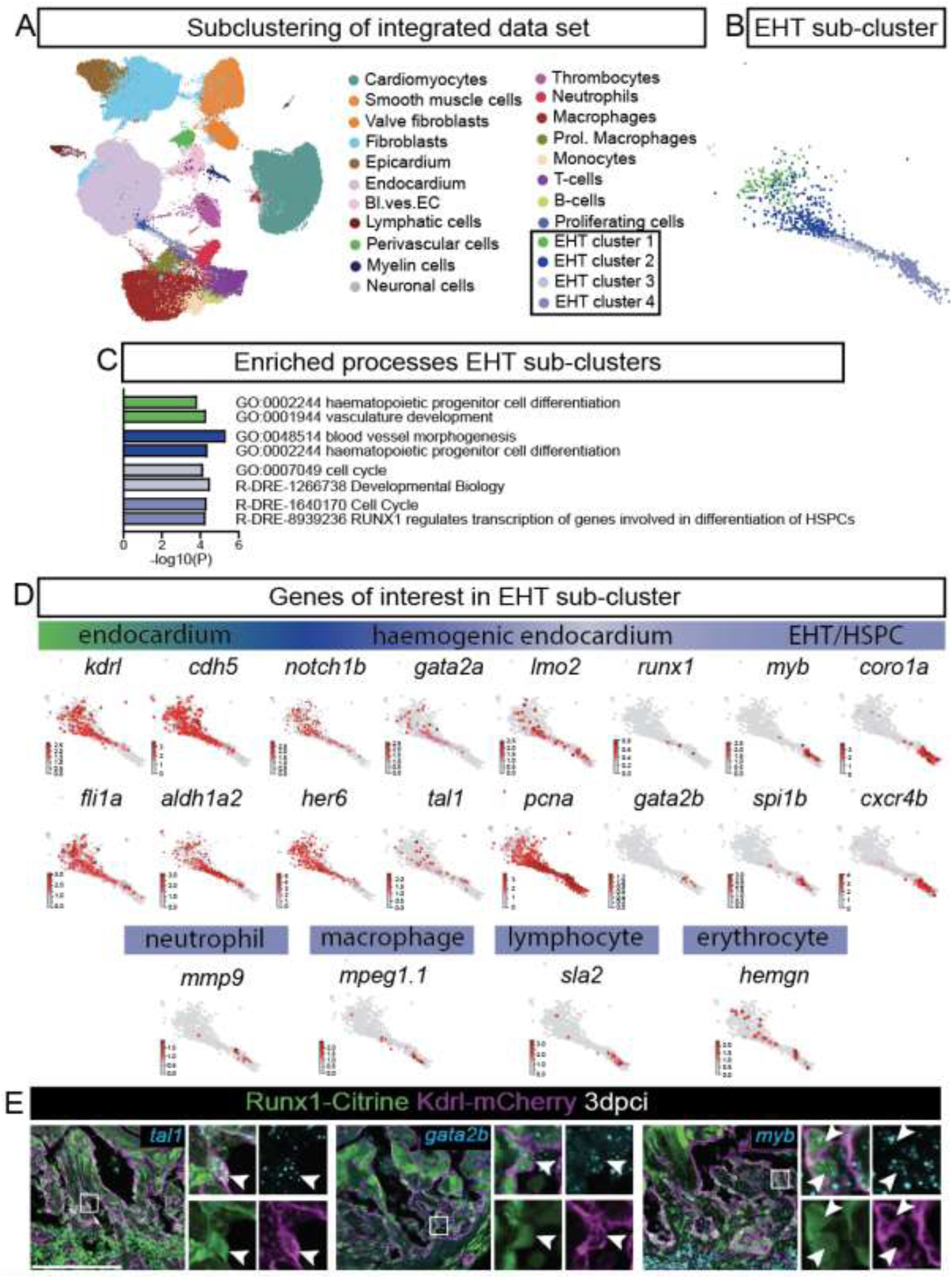
The injured endocardium gains features of endothelial-to-haematopoietic transition upon injury. A, UMAP of the integrated data set with the bridge (subclustered into 4 new clusters as EHT cluster 1-4 based on EHT progression stage) connecting the endocardium cluster with the leukocytes. B, UMAP of the 4 bridge sub-clusters described in A. C, GO terms enriched in the EHT sub-groups. D, UMAPs of key genes of interest corresponding to assigned GO terms. E, Expression of key transcription factors for EHT *gata2b*, *myb* and *tal1* at 3dpci in Runx1:Citrine positive endocardium. HSPC, haematopoietic stem and progenitor cell. Scale bar: 100µm.

Our analysis so far confirms that a subset of injured Runx1-positive endocardial cells activates an EHT gene programme. In light of our findings on the transient upregulation of a myofibroblast gene programme without much evidence of differentiation into full myofibroblasts, we questioned if the cells could similarly upregulate EHT genes as a feature of an inflammatory phenotype in endocardial cells without fully transitioning into blood lineages. Upregulation of inflammatory genes has been found in injured endocardial cells in zebrafish^45^ and endothelial cells in mouse after myocardial infarction^46–48^. Therefore, we visualised the expression of key inflammatory markers on the integrative UMAP, including *tnfa*, *nfkb* and *il1b* (Figure 3A). However, we did not observe upregulation of these inflammatory markers in the endocard ium after injury, with only marginal upregulation in the EHT clusters (Figure 3B). Specifically looking at the EHT bridging clusters, there was slightly higher expression towards the HSPC-like cluster (Figure 3C), which we have shown has lost its endocardial identity (Figure 2C-D). The published upregulation of inflammatory genes in the injured zebrafish endocardium was restricted to a very small cell population peaking at 3dpci that was identified using the Hu *et al* scRNAseq data set^34,45^. To better understand the nature of these cells in relation to our data, we also sub-clustered the endocardial cells from Hu *et al*^34^ and identified 2 small clusters with inflammatory gene profiles (Supplementary Figure 3A-B). Plotting these cells onto our integrated data set revealed that these cells were mostly distributed throughout the large endocardial cluster of our integrated data, indicating a lack of unique identity when compared to other data sets (Supplementary Figure 3C-D). However, a small subset of endocardial cells was present within the T-cell cluster of the integrated data set (Figure 1D, arrowhead in Supplementary Figure 3D), which aligns well with the reported upregulation of T-cell markers *cd74a/b* and *mhc2* genes, including *mhc2dab*, in the endocardium after injury^46^. As these cells clustered within the T-cell population and only expressed low levels of endocardial genes (Supplementary Figure 3E), we questioned if these cells could be endocardial cells differentiating into T-cells via EHT, especially as Cd4^+^ T-helper cells were previously identified directly neighbouring these cd74-positive endocardial cells^45^. Based on the upregulation of lymphoid differentiation-related genes in EHT cluster 4, (Figure 2D) and the published role of Cd74 in EHT, we then plotted *cd74a/b* and *mhc2dab* onto the EHT subclusters, which showed these genes being upregulated in HSPC-like EHT cluster 4 (Figure 3D). Therefore, in contrast to the dual temporary myofibroblast-endocardial identity we previously described, we do not find clear evidence of a temporary dual inflammatory-endocardial gene programme. However, the scRNAseq data further suggests the occurrence of EHT, including the possibility of T cells deriving from the endocardium.

**Figure 3.**
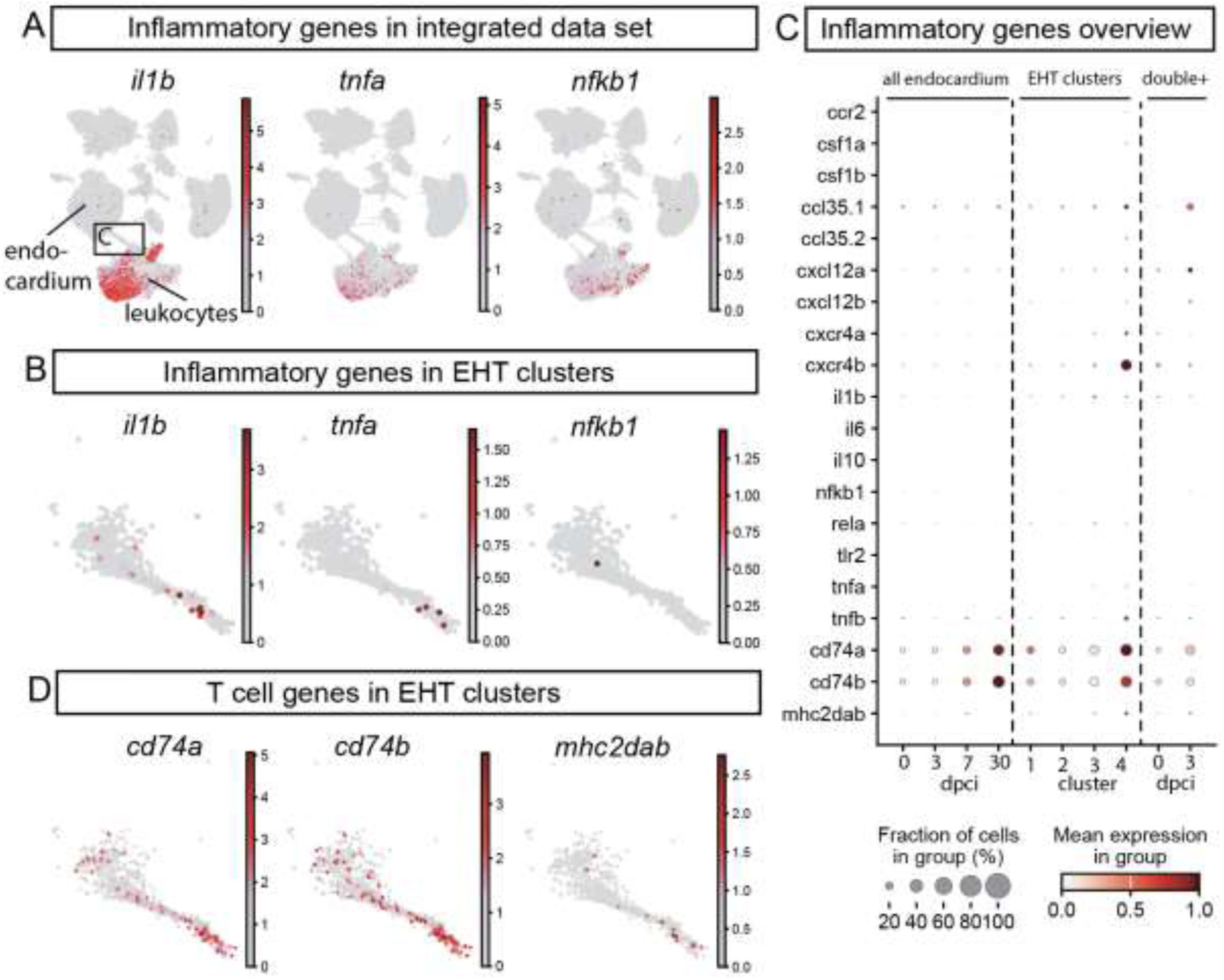
Inflammatory activation absence in response to endocardial injury. A, UMAP plots for the expression of inflammatory genes *il1b*, *tnfa* and *nfkb1* in the integrated dataset. B, UMAP of the EHT sub-clusters depicting cells expressing *il1b, tnfa* and *nfkb1*. C, Dotplot for the expression of inflammatory genes of interest in the endocardium and isolated EHT cluster or *runx1-kdrl* double positive populations after injury. D, UMAP of *cd74a*, *cd74b* and *mhc2dab* T cell gene expression in the EHT sub-clusters.

### scRNAseq lineage analysis supports the transition from endocardial to haematopoietic phenotype

In order to further investigate the suspected transition of endocardial cells towards haematopoietic lineages we performed a trajectory analysis for the cells populating the EHT bridging clusters. PAGA trajectory analysis^49^ revealed a close association among the four EHT clusters, suggesting a potential transition sequence from EHT1 to EHT4, followed by differentiation into proliferating macrophages and T cells (Figure 4A). To confirm the origin of the EHT trajectory, we utilised the unique LINNEUS approach included in Hu et al^34^. By employing Cas9, DNA scars were generated during early development, allowing lineage tracing analysis^34^. Although the scar read depth was limited to confirm the association of endothelial cells with downstream EHT clusters 1-2, there was corroborating evidence suggesting a potential transition from endocardial cells to EHT clusters 3-4 and subsequently to leukocyte populations (Figure 4B). Selecting the endocardium as the initial state for trajectory analysis in Cellrank also suggested the possibility of a gradual transition towards an HSPC-like terminal state (Figure 4C-D). This likely presents the emergence of proliferating marcophages and T cells from haematogenic endocardial cells as validated by FateID (Figure 4E-F)..

**Figure 4.**
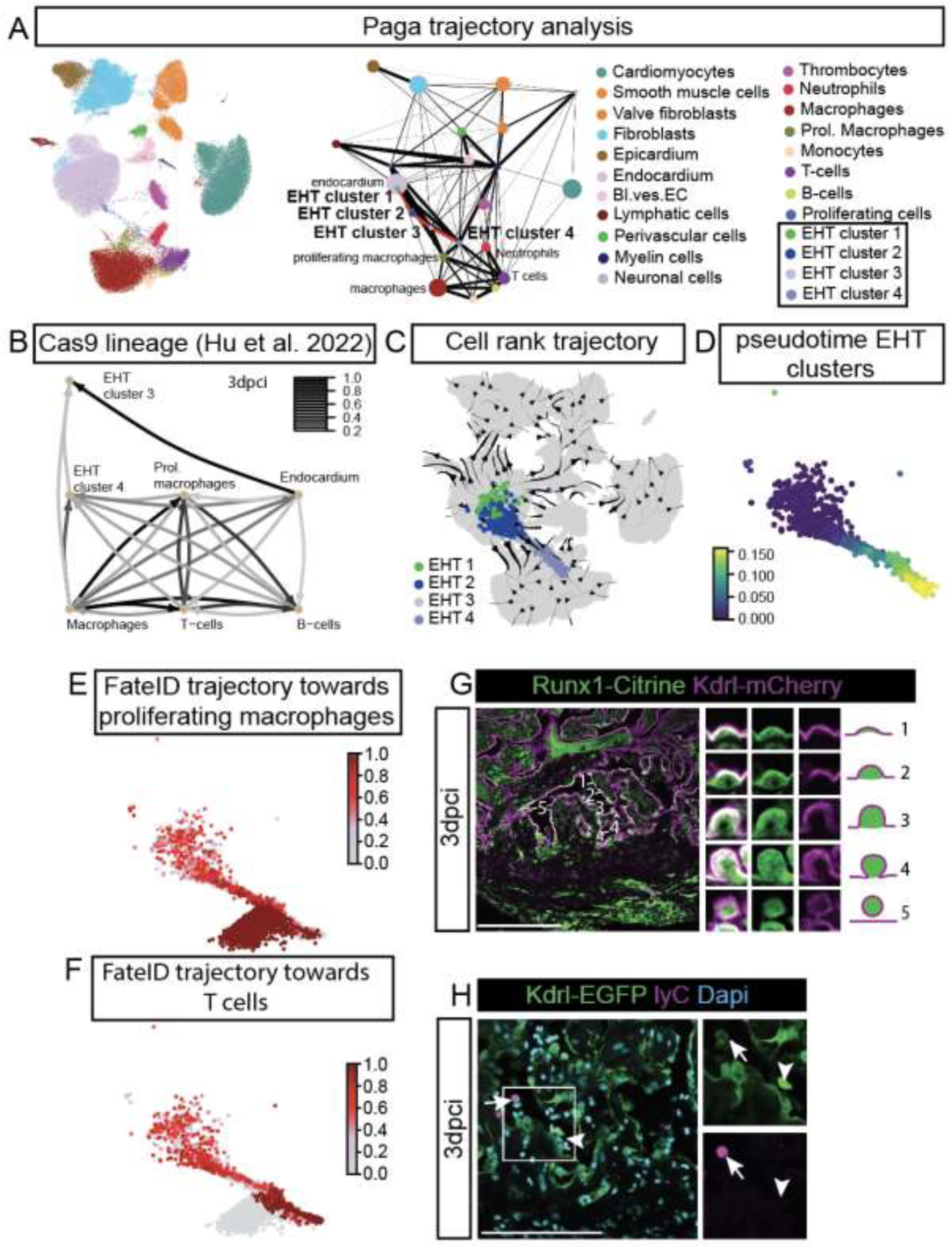
Immunostaining following heart cryoinjury reveals evidence of EHT. A, Paga trajectory analysis of transitional EHT clusters in combined data set depicts relationship of the transitional clusters from the endocardium towards white blood cell lineages. Connection thickness demonstrates a stronger cluster connectivity. B, Lineage analysis for Cas9 between EHT clusters, the endocardium and immune cells from Hu et al 2022^34^. C, Predicted cluster transition trajectories using Cellrank trajectory analysis for cell populations in the EHT clusters using the diffusion pseudotime kernelwith the endocardium cell population as the initial state. D, UMAP of transitional EHT clusters coloured by the diffusion pseudotime (DPT). E, Probabilities predicted by FateID for the transition of the EHT clusters towards a proliferating macrophage phenotype. F, Probabilities predicted using FateID for the fate transition of the EHT clusters towards T cell identity. G, Runx1:Citrine cells in the injured endocardium of Runx1:Citrine/Kdrl-mCherry zebrafish display changes in cellular morphology corresponding to cells rounding up and budding off the endocardium (morphologies 1-5). H, *kdrl-GFP*-positive circulating blood cells are found in vessels and show overlap with LyC at 3dpci. Arrowhead points to a strongly gfp-positive, *lyC-negative cell*, arrow points to a *lyC*-positive cell that appears to be losing its *gfp* expression. Scale bar: 100µm.

As the scRNAseq lineage analysis supports the transition from endocardial to a haematopoietic phenotype, we looked in more detail into the morphology of the Runx1:Citrine positive endocardial cells. It has previously been described that endocardial cells in the injury area undergo morphological changes and round up^3,4^. Considering the evidence indicating EHT occurring in the endocardium, Runx1:Citrine positive cells indeed show morphological features of cells rounding up as also seen during the phenotypic transition in the embryo, suggesting the cells could successfully transition away from the endocardial phenotype. Runx1-positive endocardial cells were observed to display different morphologies, from flat, bulging to rounded up. A small number of cells was found with no apparent connection to the rest of the endocardial sheet (Morphologies 1-5) (Figure 4G). While endocardial gene expression is turned off during EHT, it is possible to detect past gene expression due to the longer half-life of fluorescent proteins. The Tg(*kdrl:GFP*) zebrafish line has been extensively used in haemogenic endothelium studies during development because of the cytosolic labelling of endothelial cells^50^. Therefore, we next used the Tg(*kdrl:GFP*) line to examine for Kdrl-GFP-positive circulating cells on fixed sections. Small numbers of these cells could indeed be observed in the lumen near the injury site at 3dpci, with different levels of GFP-fluorescence (Figure 4H). Additionally, co-staining with a lyC antibody that labels neutrophils, showed that in some cells still weakly GFP-positive there was overlap with lyC, suggesting that these endocardial-derived cells have obtained a neutrophil identity (Figure 4H). This observation mirrors what happens in the embryonic zebrafish endocardium, where blood cells derived from haemogenic endocardium display neutrophil characteristics as marked by the positive labelling for lyz^25^. Combined, these data provide further evidence of endocardial cells gaining characteristics of haemogenic endothelium after injury of the adult heart and successfully undergoing cell fate transition.

### Live imaging and in vivo lineage tracing show the presence of endocardial-derived blood lineages

While the activation of Runx1 after injury recapitulates developmental expression and cells of the injured Runx1-positive endocardium upregulate genes involved in EHT and display the associated changes in morphology, live imaging has been essential in establishing the occurrence of this process in the embryo. To achieve this in the adult heart, we generated 75um thick live cardiac sections which were imaged for 15-24 hours, similar to a recently published approach^51^ using the Tg(*kdrl:GFP*) line which labels the endocardium with GFP and has been widely used to image the haemogenic endothelium during development. Images were taken every 10 minutes over 15 hours on 3dpci ventricle slices, the peak timepoint when EHT is indicated to occur. Examples of endothelial cells rounding up, before detaching from the endocardial surface could indeed be observed in these time-lapse movies in a similar manner to that seen during EHT in development (Figure 5A-B). The detached cells disappeared out of focal view immediately after detaching. While we were able to visualise a number of events resembling EHT, these events were rare (4/22 experiments). On one occasion, we also observed a GFP-positive cell detach from the endothelium of a vessel in the compact wall of the ventricle near the injury site. This suggests the injured vascular endothelium as another location exhibiting features of haemogenic endothelium (Figure 5C), which would fit well with the observation of upregulation of *Runx1* in the large vessels in the injury area (Supplementary Figure 1E).

**Figure 5.**
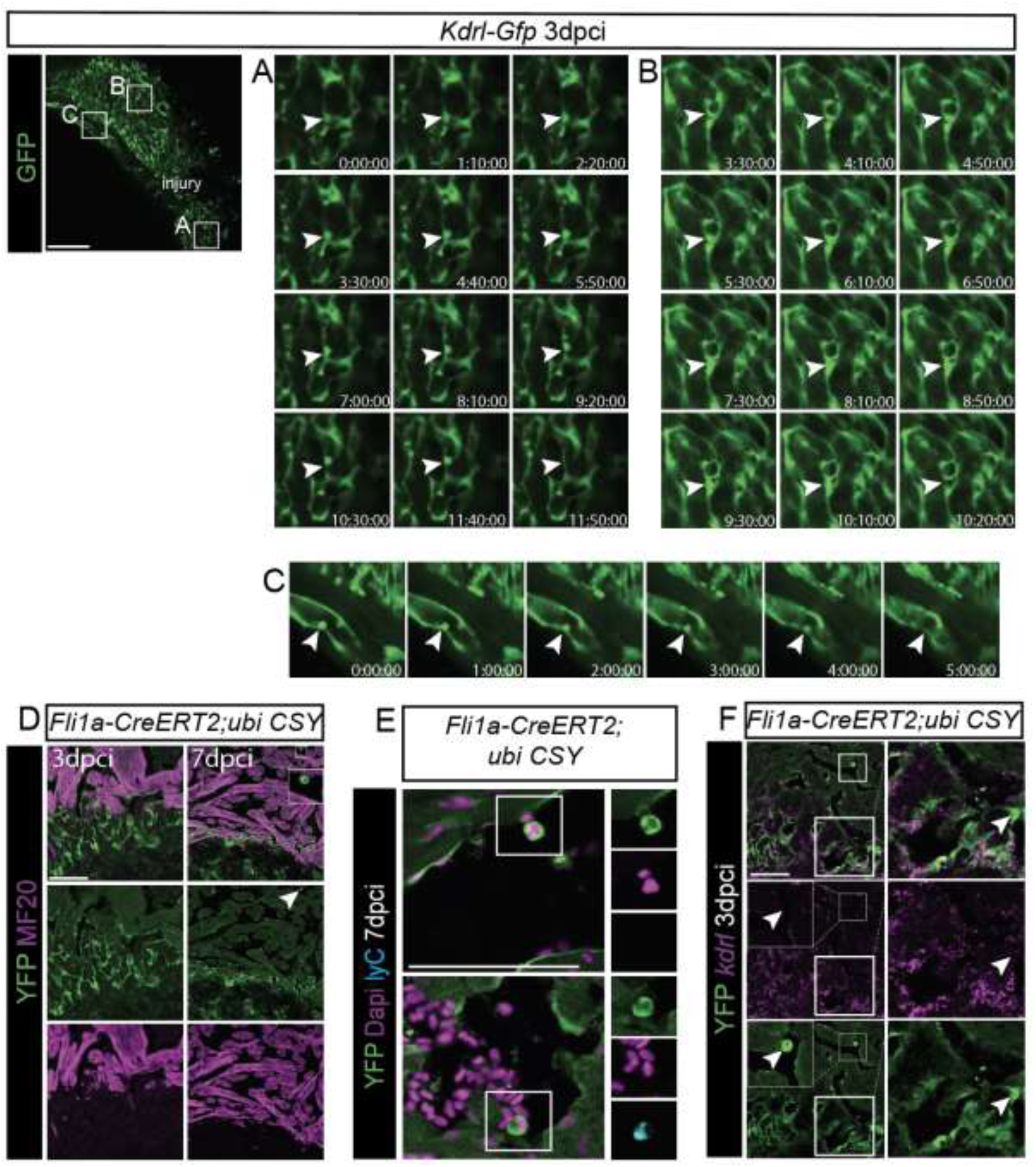
Timelapse live imaging of kdrl-GFP hearts reveals cells rounding and budding off the injured endocardium. A-B, Fluorescent imaging of live sections from injured endocardium at 3dpci over 15h shows endocardial cells became rounder, detached from the cell layer and disappeared out of view (arrowheads). C, Evidence of *kdrl*-positive cell detachment from the endothelium in a coronary vessel (arrowhead). D, Immunostaining of *YFP* and *MF20* at 3dpci and 7dpci in *Tg(fli1a-creERT2);Tg(ubiCSY)* zebrafish. The endocardium is labelled by YFP with cells in the injured area showing different morphology from the rest of the ventricle. Arrowhead at 7dpci points to YFP-positive lineage traced circulating blood cell. E, Staining of *YFP* and *lyC* reveals co-localisation of *YFP* and *lyC* expression, indicating that endocardially traced cells obtain characteristics of leukocytes. F, Immunofluorescence images for *YFP* and *kdrl* at 3dpci in *Tg(fli1a-creERT2);Tg(ubiCSY)* fish shows that while most YFP-labelled cells still express kdrl mRNA indicating that these cells are endocardial, the more rounded YFP-positive endocardial cells as well as the circulating cells do not express kdrl mRNA (arrowhead). These data indicate that rounding up endocardial cell lose their endocardial identity when they undergo EHT. This data also shows that kdrl is absent from the endocardially derived circulating cells. Scale bar: 100µm.

Next, we used the *Tg(fli1a-creERT2)* line crossed to the *Tg(ubiCSY)* line to allow for lineage analysis of the endocardium. YFP staining at 3dpci recapitulated the staining seen with the *kdrl* lines, with rounded up endocardial cells present in the injury region (Figure 5D). At 7dpci, we observed a diverse range of circulating YFP positive cells in the ventricular lumen (Figure 5D-E). Double staining for YFP and lyC indicated that while some YFP cells were lyC positive and likely neutrophils, not all were, fitting with the multilineage potential of HSPCs derived from haemogenic endothelium (Figure 5E). Importantly, while most YFP-positive endocardial cells in the wound were positive for *kdrl* mRNA, the cells with more rounded morphology did not express *kdrl* anymore, as would be expected when they undergo the transition towards a haematopoietic phenotype. In line with this, we also did not identify any *kdrl* mRNA in lineage traced YFP-positive circulating cells (Figure 5F). To further confirm that *kdrl* is not normally expressed in some circulating cells, we probed the integrated scRNAseq data for *kdrl* expression. There was no difference in the numbers of *kdrl*-positive cells before and after injury in the scRNAseq data (Supplementary Figure 4A-B). Additionally, while *kdrl* seemed to be expressed to some extent in all cell-types, including unexpectedly in cardiomyocytes and mature circulating cell populations, further analysis revealed that this could be considered background expression (Supplementary Figure 4A-C). We confirmed this on sections, where we did not observe *kdrl-mcherry* (data not shown) or *kdrl-GFP*-positive cardiomyocytes or circulating cells in the uninjured heart at any point (Supplementary Figure 4D) or any lyC-positive Kdrl-GFP-positive cells in the kidney (Supplementary Figure 4E). Additionally, if *kdrl* mRNA expressing circulating cells are present, based on the scRNAseq data, *kdrl* would be expressed in mature leukocytes, including neutrophils and macrophages and therefore likely be stable. However, the *Tg(fli1a-creERT2); Tg(ubiCSY)* lineage traced leukocytes did not express *kdrl* mRNA (Figure 5F). These results show that not only does a portion of the injured *runx1*-positive endocardium upregulate an EHT gene programme, at least a small proportion of these cells seems able to successfully transition into blood cell lineages as observed during development.

### Haemogenic endocardium is altered in absence of functional runx1

Since Runx1 is a key regulator of EHT, we next investigated how loss of Runx1 activity would influence this process, and subsequently if formation of cells with a haemogenic endocardial gene programme was impacted. The adult viable *runx1^W84X/W84X^* global mutant contains a premature truncation mutation in the Runt domain causing an almost complete loss of Runx1 activity. Crossing this line to the *TgBAC(runx1P2:Citrine);Tg(kdrl:Hsa.HRAS-mCherry)* line allowed to analyse Citrine activity via the BAC P2 promoter which serves as a readout for Kdrl-mCherry positive endocardial cells that would have expressed Runx1^52,53^. To understand any differences in EHT between the Runx1 mutant and wild-type cells in light of the previously reported difference in myofibroblast phenotype, we re-analysed the data on which the UMAP in Figure 7A from Koth et al^9^ was based. This UMAP included all *runx:citrine; kdrl-mcherry* double positive cells from wild-type uninjured and 3 dpci samples as well as 200 cells from the Runx1 mutant (Figure 6A)^9^. Among the 5 clusters previously defined, the red cluster (cluster 3) seemed to be a transition cluster located right at the boundary for transition into either a myofibroblast or haematopoietic fate. In particular, visualisation of the 5 double positive cell clusters both using a diffusion map and within the integrated UMAP highlights that after injury, the red transitionary cluster (cluster 3) exhibits a binary shift towards both cell lineages. In the wild-type condition, cluster 3 cells appeared to be in favour of the transition to the highly *myh11a* positive myofibroblasts (purple, cluster 4), while the *runx1* mutant favoured the hematopoietic cell identity as evident by the expanded cluster of highly proliferative leukocytes (orange, cluster 1) (Figure 6A-C). To better understand this phenotype in relation to all endocardial cells, next, we re-analysed 5474 cells defined as endothelial/endocardial cells from the Koth data set^9^ and reassigned these cells into 13 distinct clusters (Supplementary Figure 5A-B, Figure 6D). On the UMAP, 2 injury-responsive clusters appear in the wild-type at 3dpci, one characterised by the myofibroblast marker *tagln* (cluster 11) and the other by high *pcna* expression levels with cells in S and G2M phase (cluster 8). In contrast, the myofibroblast cluster (Cluster 11) is absent in the *runx1* mutant, but there is an increase in *pcna* positive cells (Figure 6D-E, Supplementary Figure 5C). To determine if this *pcna* positive cluster serves as a progenitor pool emerging from the endocardial cluster and as a progenitor pool for a downstream cell type, we performed RNA velocity analysis capturing full transcriptional dynamics (Figure 6E). This showed that the *pcna*-positive cluster transitions away from the endocardial cells, suggesting fate transition towards another cell phenotype. Plotting 3dpci *runx1* mutant and wild-type *pcna* on the integrated UMAP shows that *pcna* expression reflects the pattern of the *runx1:citrine, kdrl-mcherry* double positive cells within the leukocyte cluster and suggests a higher occurrence of EHT in the mutant (Figure 6F). Therefore, Runx1 may be responsible for regulating the balance between obtaining a myofibroblast versus EHT cell fate.

**Figure 6.**
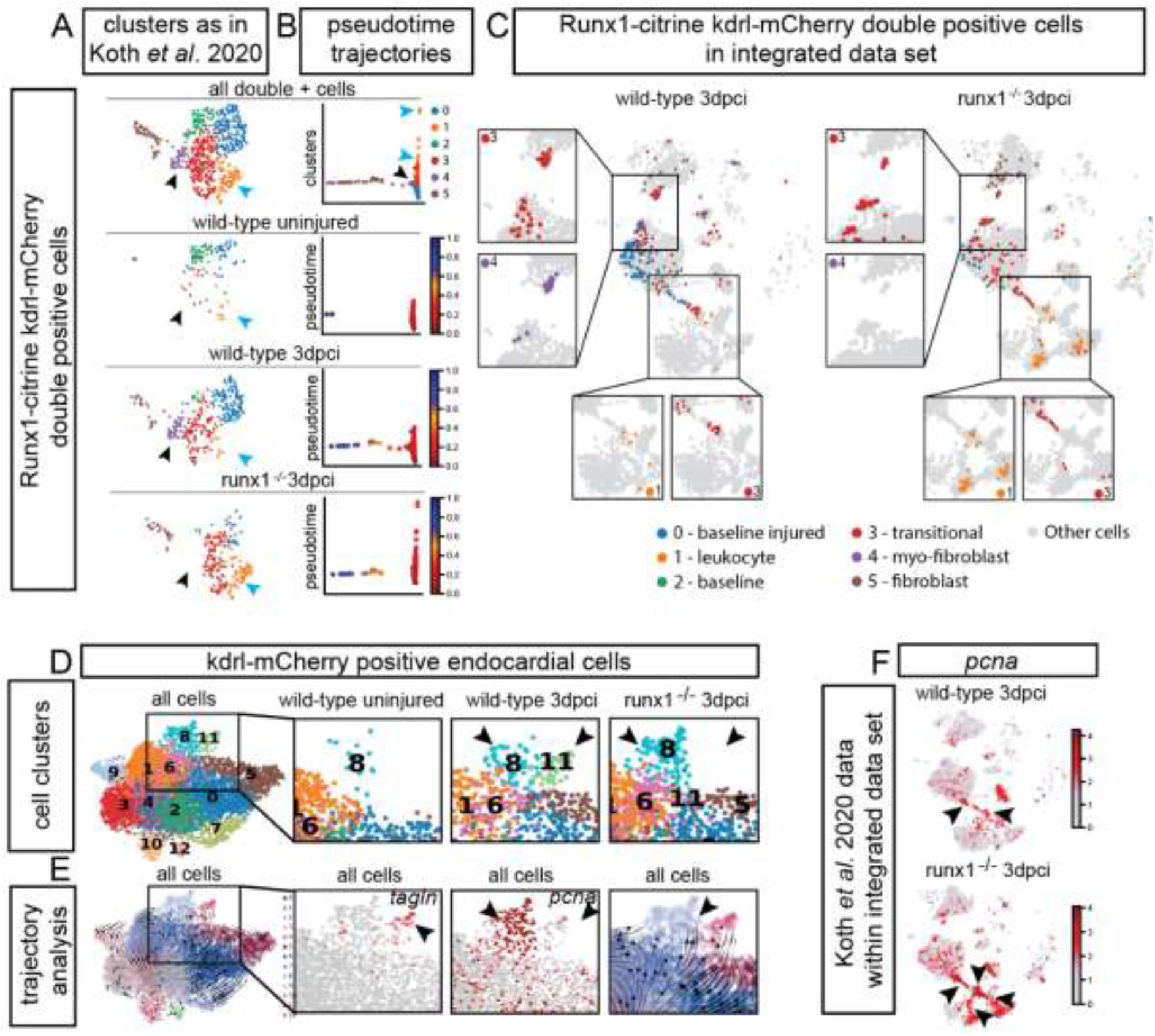
Interrogation of Koth et al 2020 data set clustering in integrated data set. A, UMAP of *runx1-kdrl* double positive cells from Koth et al 2020^9^ showing the previously defined 6 sub-clusters. B, Diffusion maps of double positive *runx1-kdrl* cells coloured by the diffusion pseudotime initiated from sub-cluster 0, across different clusters in wild type uninjured control and 3dpci in wild type and Runx1 mutants. C, *runx1-kdrl* double positive cells from Koth et al^9^ dataset localisation in integrated data cell clusters in wild type and Runx1 KO following cryoinjury. D, Clustering of kdrl positive cells in wild-type control, wild type 3dpci and runx1 KO at 3dpci shows expansion of endo-sub-cluster 8 with injury and absence of endo-sub-cluster 11 in the mutant. E, RNA velocity analysis of kdrl expressing endocardial cells in global cell population and highlights on tagln and pcna expression (inserts). F, UMAP displaying localisation of *pcna* expressing cells in wild-type and runx1 mutants at 3dpci from ^9^ in integrated data set clusters.

**Figure 7.**
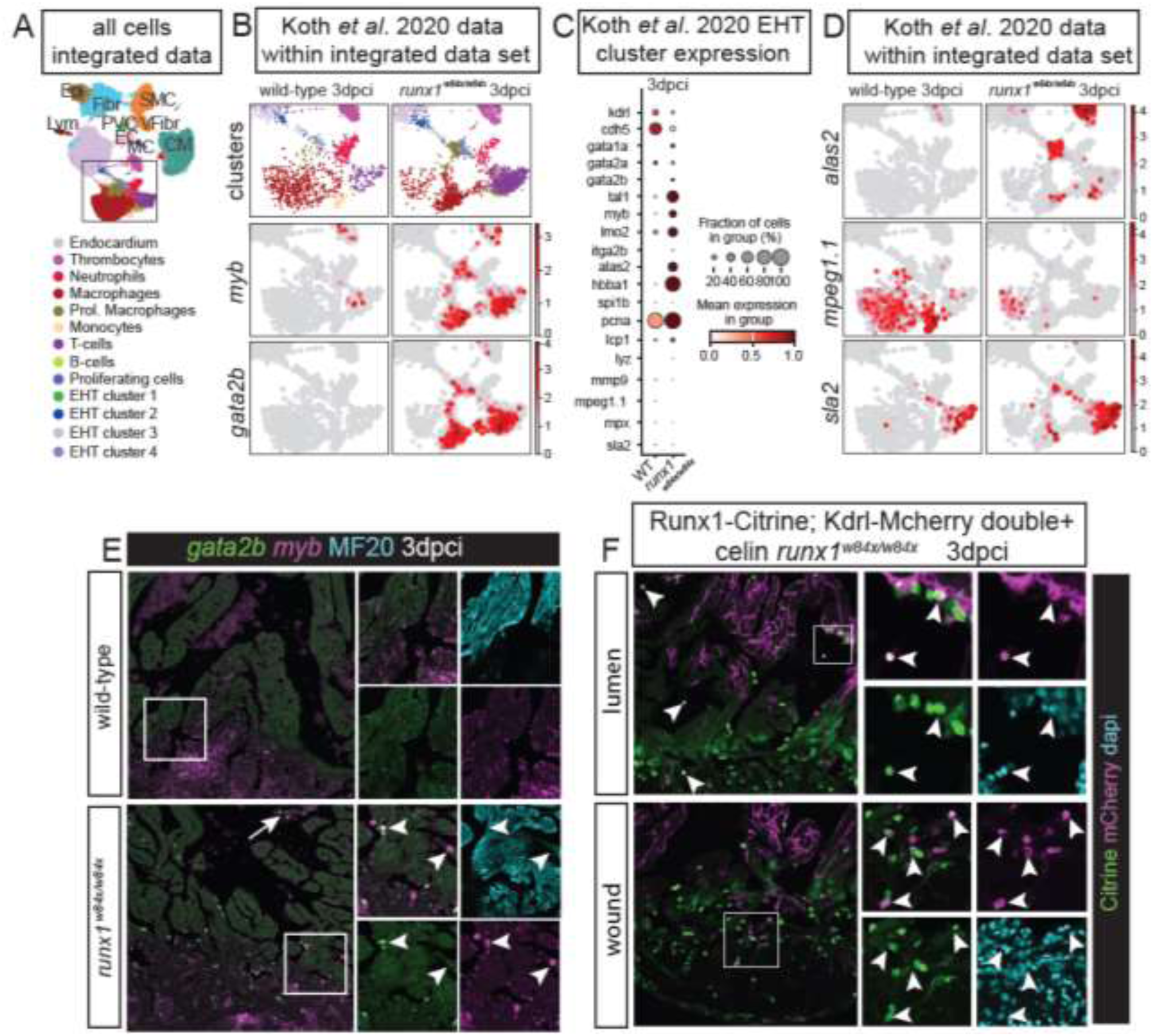
Cellular lineage shift observed with *Runx1* absence. A, UMAP of all integrated datasets with focus on blood lineage clusters arising from the large endocardial cluster as displayed in ^9^. CM, cardiomyocytes; EC, endothelial cells; Epi, epicardial cells; Fibr, fibroblasts; Lym, lymphatic cells; MC, myelin cells; PVC, perivascular cells; SMC, smooth muscle cells; VFibr, valve fibroblasts. B, UMAP displaying the Koth *et al* dataset^9^ for wild-type and runx1 mutant hearts 3dpci extracted from the integrated landscape. Focus on *myb* and *gata2b* displays increase in expression of both genes in Runx1^w84x/w84x^. C, Dotplot for expression levels of key genes in EHT cluster 3dpci in Koth *et al* dataset^9^ for wild-type and runx1 mutant. D, UMAP of Koth et al ^9^ data location in the integrated data set UMAP clusters for *alas2*, *mpeg1.1* and *sla2* in both wild-type and mutant *Runx1* conditions. E, Immunohistochemical staining for gata2b, myb and MF20 expression in wild-type and runx1 mutant. Arrowheads point to double positive cells for *gata2b* and *myb* in the endocardium. F, Expression of *Runx1:Citrine* and *Kdrl-mCherry* in *Runx1* mutant heart 3dpci. Double positive cells are visible in both the injury site and lumen close to the wound (arrowheads). Scale bar 100µm.

### Absence of Runx1 alters cell identity programmes

Following the observation that in the absence of functional Runx1 there is a switch from a myofibroblast gene programme towards a leukocyte gene programme, we looked into how these EHT-associated cells distributed among the different leukocyte compartments in the presence or absence of Runx1 (Figure 7A-B). Interrogation of the Koth data set in the integrated data at 3dpci in the *runx1^W84X/W84X^* mutant displayed an expansion of the macrophage and T-cell compartments when compared to the injured wild-type (Figure 7B). Specifically, mutation of functional Runx1 promoted an increase in highly proliferative cells expressing *myb* and *gata2b*, two transcription factors known for their role in regulating proliferation and cell differentiation decisions in the haemogenic endothelium and HSPCs (Figure 7B-C)^54–56^. These highly *myb* and *gata2b* expressing blood cells were clearly visible on 3dpci sections as also previously reported (Figure 7D)^9^. We subsequently tested whether this had a downstream effect on the induction of haematopoietic differentiation programmes and the populations of myeloid and lymphoid cells derivatives. Comparison of the expression of haematopoietic regulators and EHT markers in the bridging EHT clusters (Figure 2B) in wild-type and mutant showed a strong increase of EHT-related genes as well as proliferation and erythropoiesis genes in the *runx1* mutants (Figure 7C). We have previously observed that the integrated data set did not show significant evidence of erythrocyte differentiation as marked by *tal1* expression occurring in the HSPC-like cluster (Figure 2D). Deletion of *runx1*, however, seemed to promote erythrocyte differentiation, albeit that there was evidence of overall increased erythrocyte lysis in the mutant that could have biased this finding. T cell differentiation was also increased in the mutant as evident by the increase in *alas2* and *sla2* positive cells (Figure 7E). Conversely, macrophage differentiation was reduced as shown by the reduced numbers of *mpeg1.1*-positive cells in relation to wild type in the Koth data in the integrated dataset.

This preference in cellular transition in the mutants was also observed on tissue sections where smaller and immature looking Runx1:Citrine; Kdrl-mCherry positive circulating cells were present at the injury site in the *runx1* mutant, both within the lumen as well as within the wound (Figure 7F). Together, these results further support our observation that haemogenic endocardium is altered in the absence of *runx1.* Thus, Runx1 may be involved in the regulation of the balance of different EHT branches or could be responsible for inducing macrophage formation while alternative regulators may be present for the induction of erythroid and T cell programmes.

## Discussion

During development, endocardial cells mainly provide the inner lining of the heart but also contribute to the formation of other cardiac cell populations including fibroblasts of the heart valves, the coronary vasculature and blood cells^57^. Cardiac injury re-activates endocardial developmental processes and here we show that upregulation of Runx1 in the injured endocardium recapitulates expression observed in the embryo during development and differentially regulates gene expression programmes governing endocardial identity following injury.

EHT was initially reported to occur in the embryonic mouse heart^26,27,58^. However, the occurrence of EHT has since been questioned as endocardial lineage analysis using Cre lines did not identify endocardial derived blood cell lineages^59,60^. Lately, there has been renewed interest in this process due to possible discrepancies in reported findings of Cre expression systems. Multiple groups report the embryonic endocardium behaving similar to haemogenic endothelium and contributing to erythroid, platelet and myeloid lineages in mouse^26,58,61^ as well as zebrafish^25,28^. Our observation of Runx1 expression, a known marker of haemogenic endothelium^62^, in the developing endocardium (Figure 1A) aligns with recent reports of EHT and transient haematopoiesis occurring in the zebrafish endocardium during development^28,61^.While one study reports 24hpf as the peak of EHT ^25^, we only see the initiation of Runx1expression from around 2dpf. The peak of Runx1 expression we observed at 5dpf fits better with more recent reporting of the first Runx1-positive cells attached to the endocardium at 56hpf and EHT peaking at 72hpf^28^. While we have not further investigated the formation of the embryonic haemogenic endocardium, this suggests multiple waves of EHT occurring during development, with Runx1 involvement in the later wave. Multiple scRNAseq studies have highlighted the heterogeneity of EHT for the generation of the various haematopoietic cells^36,37,40,63,64^. Therefore, it could be possible that two waves of induction of haemogenic endocardium occur in the embryo which contribute to different blood lineages. The first wave of EHT at 1dpf seems likely to mostly give rise to neutrophils^25^, while other cell types may be derived in a subsequent induction of haemogenic endocardium around 3-5dpf.

In addition to the previously published transition into a myofibroblast phenotype^9^, we now find evidence of another sub-population of Runx1-expressing endocardial cells in the adult heart after injury that transition towards a haematopoietic fate. The expression of Runx1 is key for this lineage decision, as in absence of functional Runx1, the cells shift from the myofibroblast lineage towards the haematopoietic identity. A similar role for Runx1 during cell identity transition has also been described in other organs, where Runx1 determines the balance between proliferation and differentiation, with control over cell-type transitions towards myofibroblast^14^ and haematopoietic lineages^65^. However, how some endocardial cells are primed to make those decisions and what triggers and regulates these gene expression changes is not known. A transient identity shift towards a smooth muscle cell-like phenotype, without loss of the endocardial identity, could be key for successful scar degradation and regeneration. Likewise, the upregulation of an EHT gene programme in the endocardium might largely be transient, with only a subset of cells actively undergoing full transition to HSPCs as similarly to our lineage tracing of myofibroblasts, we found only few circulating haematopoietic cells with an endocardial origin. The activation of EHT seems to have a positive impact on regeneration as EHT is enhanced in the faster regenerating *runx1* mutant^9^, highlighting that Runx1 expression after cardiac injury is suboptimal for tissue repair^18,19^ which possibly facilitates scarring of the heart.

Injured cardiac endothelial and endocardial cells have previously been shown to also upregulate inflammatory gene programs^46–48,66^ and cytokine signalling from the endocardium is known for its role in regeneration, specifically for regulating EMT and inducing myocardial proliferation^5,67.^ Therefore, expression of Runx1 could also be important for activation of a transient inflammatory state in response to endocardial injury. However, we identified no clear activation of inflammation in the injured Runx1-positive endocardium. This lack of upregulation of a broad inflammatory gene programme in response to injury could be a critical difference to the mammalian response after injury in terms of regenerative capacity. Some injured endocardial cells have recently been found to upregulate antigen presentation genes, which have been suggested to communicate to infiltrating adaptive immune Cd4+ T helper cells^45^. Tracing these antigen presenting endocardial cells in our integrated data set suggests that these cells might part of the population of cells undergoing EHT that is differentiating towards a T cell identity. This would fit well with that the T helper cells were always found near the antigen presenting endocardium^45^.

The haemogenic endocardium after injury seems to give rise to multiple blood lineages, in particular macrophage and lymphocyte lineages, but with some neutrophil and erythrocyte cells. We have previously shown that the *runx1* mutant has abnormal blood cells in the heart, expressing high levels of *myb* and *gata2b*^9^. Recent findings on blood cell populations in the *runx1* mutant^68^ highlight that these haematopoietic populations are also present in absence of injury. Since *runx1* is known as a key regulator during definitive haematopoiesis, it is unclear how some *runx1* mutants survive during early development in absence of definitive haematopoiesis in the dorsal aorta, and still develop multilineage blood populations. The endocardium could hence be the source of these blood cells during development for which Runx1 function during EHT is dispensable. This would support our data, showing that in absence of *runx1*, a shift towards EHT versus myofibroblast identity occurs after injury. Further evidence shows that endocardial-specific deletion of Runx1 does not affect the production of endocardial-derived macrophages from multipotent hematopoietic progenitors during mouse embryonic development, revealing that endocardial-derived macrophages are not Runx1-dependent and hence, not definitive in nature^26^. We, however, observed that deletion of *Runx1* resulted in the increase of erythrocyte and T cell populations but reduced the cluster size for mpeg1.1 positive macrophages (Figure 7D) which does not fully align with the embryonic mouse data. Our observation for the reduction of the macrophage cluster and shift towards T-cell and erythrocyte phenotypes thus could imply the possible presence of distinct haematopoietic pathways present in the adult zebrafish heart. Further work is needed however to identify the specific mechanisms via which these varied responses arise.

There is evidence that leukocytes with an endocardial origin may have unique functions. In particular, EHT-derived macrophages from the developing mouse valves were shown to persist from E10.5 to adulthood and exhibit a distinct predominantly M2 phenotype displaying higher levels of phagocytosis^26^. This suggests that macrophages (and other leukocyte populations) derived from the injured endocardium could be distinct from macrophages from other sources and have different functions after injury. As EHT is increased in the Runx1 mutant and this mutant has enhanced regeneration, leukocytes derived from the EHT process could have specific positive functions during regeneration. Specifically, higher levels of phagocytosis by macrophages after injury could increase scar degradation and speed up regeneration, which aligns with the reduced levels of scarring we previously observed^9^. However, the function and contribution to heart regeneration of endocardial-derived leukocytes requires further investigation as we identified the presence of both lyC positive and negative circulating cells which is suggestive of endocardial contribution to different leukocyte compartments. Our study combines extensive bioinformatic and *in vivo* analysis to show that EHT mechanisms are reactivated following injury and that Runx1 is involved in this process. This is an exciting finding that will need further validation in follow up studies as it could play a key role during the regenerative process. In particular, the extent to which endocardial cells undergo the full transition towards leukocyte populations after injury needs further investigation as well as the role these unique blood populations could play during successful regeneration. Additionally, further research into the role of Runx1 as regulator of the balance between cardiac fibrosis and successful regeneration will increase our understanding of the cardiac injury response and provide candidate targets for therapeutic modulation in the injured heart^69^.

## Author contributions

Conceptualisation: X.W and M.T.M.M

Methodology: X.W and M.T.M.M

Data collection: all authors

Data analysis and curation: J.Y, I.L, Y.K, A.K., X.W. and M.T.M.M

Writing-original draft: I.L and M.T.M.M.

Writing-editing: J.Y, I.L, Y.K, X.W. and M.T.M.M

Reviewing: all authors

Visualisation: J.Y, I.L, Y.K, A.K., X.W. and M.T.M.M

Supervision: X.W and M.T.M.M.

## Acknowledgements

We thank Henry Roehl for providing the *Tg(Fli1a:ERT2CreERT2)* line. Oxford’s BMS teams for zebrafish husbandry support. This work was supported by a BHF non-clinical PhD fellowship FS/14/73/31107 (A.K and M.T.M.M.) and by the European Research Council (ERC) under the European Union’s Horizon 2020 research and innovation program (grant agreement n° 715895, CAVEHEART, ERC-2016-STG, M.T.M.M.). This work was also supported by the BHF Centre of Regenerative Medicine (RM/13/3/30159), the BHF Centre of Research Excellence Oxford (RE/13/1/30181) and the National Natural Science Foundation of China (82470478, X.W). The imaging facility used for this study was supported by the MRC via the WIMM Strategic Alliance (G0902418).

## Methods

### Zebrafish strains and husbandry

All experiments were carried out under appropriate Home Office licenses and in compliance with the revised Animals (Scientific Procedures) Act 1986 in the UK and Directive 2010/63/EU in Europe, and all have been approved by Oxford’s central Committee on Animal Care and Ethical Review (ACER). Adult wild-type (WT) from the KCL strain, *Tg(kdrl:Hsa.HRAS-mCherry)*^70^, *runx1^W84X^* ^53^, *TgBAC(runx1P2:Citrine)*^29^, Tg(kdrl:EGFP)^s843 50^, *Tg(Fli1a:ERT2CreERT2)* obtained from Henry Roehl (University of Sheffield, UK), and *Tg(ubb:LOXP-AmCyan-LOXP-ZsYellow)^fb^*^5^ referred to as *ubi:CSY* ^71^, were maintained in the KCL background and housed in a Techniplast aquarium system [28 °C, 14/10 hours light/dark cycle, fed 3x daily with dry food and brine shrimp]. All double transgenic lines on wild-type or mutant background were generated by natural mating. Zebrafish embryos were kept in petri dishes with E3 embryo medium in incubator at 28°C^72^ until required for fixation.

### Cardiac surgery

Zebrafish cryoinjury of the ventricle was performed as previously reported^73,74^. Briefly, prior to all surgical operations, fish were anaesthetised in MS222 (Sigma). A small incision was made through the thorax and the pericardium using forceps and spring scissors. The abdomen was gently squeezed to expose the ventricle and tissue paper was used to dry the heart. A cryo-probe with a copper filament was cooled in liquid nitrogen and placed on the ventricle surface until thawing was observed. Body wall incisions were not sutured, and after surgery, fish were returned to water and stimulated to breathe by pipetting water over the gills until swimming was restored. All operated fish were kept in individual tanks for the first week after surgery, then fish were combined in larger tanks.

### Tissue Processing

Hearts were extracted and isolated in PBS. Hearts were inspected, cleaned and then fixed with 4% Paraformaldehyde (PFA) overnight (O/N) at room temperature. Samples were dehydrated into Ethanol (EtOH) at 70%, 80%, 90%, 96% for 2 hours each step and 2×100% for 1 hour each step, followed by a 100% 1-butanol step overnight. The samples were then transferred to paraffin wax (Paraplast Plus, Sigma-Aldrich P3683) at 65°C. Paraffin was refreshed 2x with each step at least 2 hours, prior to mounting in a sectioning mould. 10µm sections were cut using a Leica microtome and section ribbons were stored on black cardboard in shallow stackable plastic trays. Individual sections, evenly distributed throughout the heart e.g. 1 in 6 sections, were selected and mounted on superfrost plus glass slides for immunohistochemistry and RNA labeling (RNAscope).

### Embryo Processing

Zebrafish embryos were fixed in 4% PFA for 45 minutes at room temperature in 1.5ml tubes while very gently rocking on a nutator, then washed in PBTr [PBS with 0.5% Triton (Triton X-100, T8787 Sigma-Aldrich) three times for 5 minutes.

### Immunohistochemistry

Fluorescent immunohistochemistry on sections was performed as previously described^75^. For de-waxing, slides were taken through 2x Xylene 5minutes, 2x 1 minute in 100% EtOH 2 minute, 1x 96%, 90%, 80% and 70% EtOH, and a final rinse in PBS prior to subsequent staining. De-waxed and rehydrated sections were heated up and then pressure cooked for 4 minutes in antigen unmasking solution (H-3300, Vector Laboratories Inc). Once cooled, sections were placed in PBS before drawing a ring (ImmEdge pen, Vector Laboratories) around the sections. Slides were placed into staining trays providing humidity and blocked using TNB (0.5% TSA blocking reagent, 0.15 M NaCl, 0.1M TRIS-HCL, pH 7.5, NEL702001KT, Perkin Elmer) for 30minutes at RT. Blocking agent was removed and primary antibody in TNB was added and incubated O/N at RT. Slides were then washed 3x 5 minutes in PBS before the secondary antibody (Alexa range, Invitrogen) at 1:200 dilution in TNB was added for 2 hours at RTSlides were mounted in Mowiol 4-88 (Applichem) and slides incubated at 37°C O/N in the dark. The following primary antibodies were used: chicken polyclonal against Green Fluorescent Protein (GFP, 1:200, Aves Lab, GFP-1020), mouse monoclonal against mCherry (clone 1C51, 1:200, Abcam, ab125096), Proliferating Cell Nuclear Antigen (PCNA, clone PC10, 1:200, Dako Cytomation, M0879), Myosin Heavy Chain (MF20, 1:50, HSHB AB-2147781), cardiac Troponin-T (cTnT, 1:500, Abcam, ab33589) and rabbit polyclonal against Lysozyme (LyC, 1:200, Anaspec, AS-55633), smooth muscle Myosin heavy chain 11 (Myh11, 1:200, Abcam, ab125884). For double labelling with RNAscope probes, RNAscope was performed first and then processed for immunohistochemistry as described above, starting from the blocking step.

Fluorescent immunohistochemistry on embryos was performed immediately after fixation and 3 PBTr washes, by blocking 1 hour with 5% Goat serum in PBTr, followed by primary antibody incubation on nutator at 4°C O/N in the dark, followed by 6 x 15 minutes washes with PBTr at RT, followed by secondary antibody incubation on nutator at 4°C O/N in the dark, followed by 6 x 15 minutes washes with PBTr at RT and mounting on coverslips with VECTASHIELD® Antifade Mounting Medium (H-1000-10).

### RNAscope In Situ Hybridisation

RNAscope® (Advanced Cell Diagnostics, Hayward, CA)^76^ was performed on 10µm thick paraffin sections, processed as described above. Sections were baked at 60°C for one hour before deparaffinisation using 2x 5 minutes Xylene steps followed by 2x 2 minutes 100% EtOH. The slides were air-dried followed by incubation in RNAscope Hydrogen Peroxide (H_2_0_2_) for 15 minutes before washing in MilliQ. The slides were then boiled at 98-102C for 15 minutes in 1x RNAscope® Target Retrieval solution, placed in 100% EtOH for 3 minutes and air-dried. The sections were the incubated with RNAScope Protease III in a Hybez oven at 40°C for 15 minutes, washed in MilliQ 2x 2 minutes, followed by incubation with the different RNAScope probes for 2 hours at 40°C. The RNAscope Multiplex Fluorescent Detection Reagents v2 and the TSA Plus Cyanine 3 and 5 fluorophore (Perkin Elmer, NEL744001KT) were applied according to the manufacturer’s instructions. The slides were further processed for immunohistochemistry or mounted in Mowiol 4-88. Advanced Cell Diagnostics designed the probes. Probes used were Dr-myb-C3 (558291-C3) and Dr-gata2b-C2 (551191-C2).

### Slice cultures

Hearts were isolated at 3dpci and placed in chilled PBS on ice. Excess blood and tissue was removed from the outside of the hearts before hearts were gently dried with tissue paper. The hearts were then placed in the middle of a square mould and embedded in 4% LMP-agarose in PBS. The blocks were then sectioned on a vibratome (Leica, VT1000S) with settings of 1.1mm/s at 600Hz with a blade from Astonics, INC (89-0015) into 75μm slices and collected in a bath of chilled PBS. Slices were captured using a pair of fine-tipped forceps and placed into a dish containing chilled PBS on ice. Next, a welled coverslip was placed onto a hot plate at 42 °C and molten LMP agarose was added to the wells before placing the tissue slices on top of the agarose with a pair of fine-tipped forceps. Excess agarose was from the well, leaving a thin layer on the bottom. The coverslip was then removed from the hot plate allowing the agarose to solidify. The wells were then covered with culture medium (DMEM, 10% fetal bovine serum, 1% MEM-NEAA, antibiotics/anti-contaminants (100U/ml penicillin, 100ug/ml streptomycin, primocin) and 50µM beta-mercaptoethanol) and a microscope slide was placed over the top. Any excess medium was removed from the sides of the slide before imaging.

### Image acquisition and data analysis

Paraffin section images were acquired using either a Zeiss LSM880 or Olympus FV3000 confocal. Vibratome sections were imaged on a Zeiss LSM 880 Airy Scan, with images taken at intervals for periods as described. Images were processed in FIJI/ ImageJ to generate a magenta/green/cyan/grey color scheme.

### Single-cell RNA-Seq datasets

The scRNA-Seq datasets used in the study include: 1) GSE138181 (Koth)^9^; 2) GSE153170 (Bakker)^30^; 3) GSE172511 (Sun)^31^; 4) GSE159032 and GSE158919 (Hu)^34^; 5) GSE188511 (Kapuria)^32^; and 6) GSE145980 (Ma)^33^. A complete summary of samples can be found in Zebrafish_FinalTable.xlsx.

### Single cell sequencing data analysis

The processed scRNA-Seq data from the six studies were obtained from the Gene Expression Omnibus (GEO)^9,30–34^. Preprocessing of the scRNA-seq dataset was performed using Scanpy (v.1.9.3)^77^. After preliminary quality control (Supplementary Table 1), the remaining 289,465 cells were normalised to a final count of 1e^4^ and log-transformed. A total of 1360 highly variable genes were selected with ‘min_mean=0.02, max_mean=3, min_disp=0.3’. The datasets were then integrated using the Harmony algorithm with the top 50 PCA components^78^. An additional 10081 cells with a low number of gene contents were removed. Cell types were defined according to the annotation from Hu et al. ^34^. Each non-Hu cell was assigned to the 15 closest neighbouring cells from the Hu data based on Euclidean distance calculated using the top 50 principal components and the cell type was assigned based on the most frequently occurring type.

To investigate the EHT clusters, Leiden clustering was performed on all cells with the resolution set to 0.8, resulting in 24 clusters, referred to as Leiden_v1. This was further refined by increasing the resolution to 2.0, which yielded 39 clusters, referred to as Leiden_v2 (Supplementary Figure 6A-B). Leiden_v2 was used in Figure 2, 4A-F and Figure 7. To further analyse the EHT, Cluster 25 in Leiden_v2 was subclustered into 4 EHT clusters using a restricted clustering approach with the ‘restrict_to’ parameter and a resolution of 0.3. T cells or macrophages as per Hu’s cell type annotation within cluster 15 of Leiden_v1 were further identified as proliferating T cells and proliferating macrophages, respectively. Leiden_v1 was used in Figure 4E-F. The rank-gene-groups function in Scanpy^77^ was used to identify marker genes for each subcluster with the method set to ‘t-test’. UMAPs and dotplots were generated using Scanpy^77^.

To further investigate the inflammatory response in the endocardium after heart injury, the log-transformed normalized counts of the endocardial cells from Hu et al’s data were extracted and used to re-calculate the UMAP using the same parameters as mentioned above. Leiden clustering was performed with the resolution set to 1, which yielded 17 endocardial subclusters. 2 clusters (cluster 13, 14) expressing inflammatory genes (Supplementary Figure 3) were annotated as inflammatory endocardial cells^45^.

### Lineage analysis

Scar lineage tracing information was obtained from the GEO database (GSE159032). Lineage tree construction was performed using the R script Iterative_tree_building. R in the original manuscript, excluding the DMSO/IWR/morphine treatment samples^34^. Nodes with fewer than two cells were removed, and conditional transition probabilities were calculated as described in the paper. The transition network was visualized using Igraph v.1.5.0^79^ in R with arrows indicating probabilities greater than 0.2.

### Trajectory analysis

PAGA^49^ (Partition-based graph abstraction) plots were calculated and visualised using Scanpy^77^ with default parameters, displaying lines for connectivity >0.0001 and dot sizes scaled to population size. Cellrank v.2.0.0^80^ was used to construct the transition matrix using the diffusion pseudotime kernel, with one cell in EHT cluster 1 selected as the root cell (Supplementary Figure 6C). FateID v.0.2.2^81^ was used to estimate the fate probabilities from the EHT clusters to T cells or proliferating macrophages.

### Background contamination analysis

SoupX v.1.6.2^82^ was used to estimate the cell-free ambient RNA contamination probability using the raw count matrix generated by Cellranger v.6.1.2^83^ with default parameters.

### Data availability

Scripts used in this manuscript have been deposited at https://github.com/JasonJunYing/EHT. Due to the size of the raw data, it can be provided upon requests.

## Supplementary Figures

**Supplementary Figure 1.**
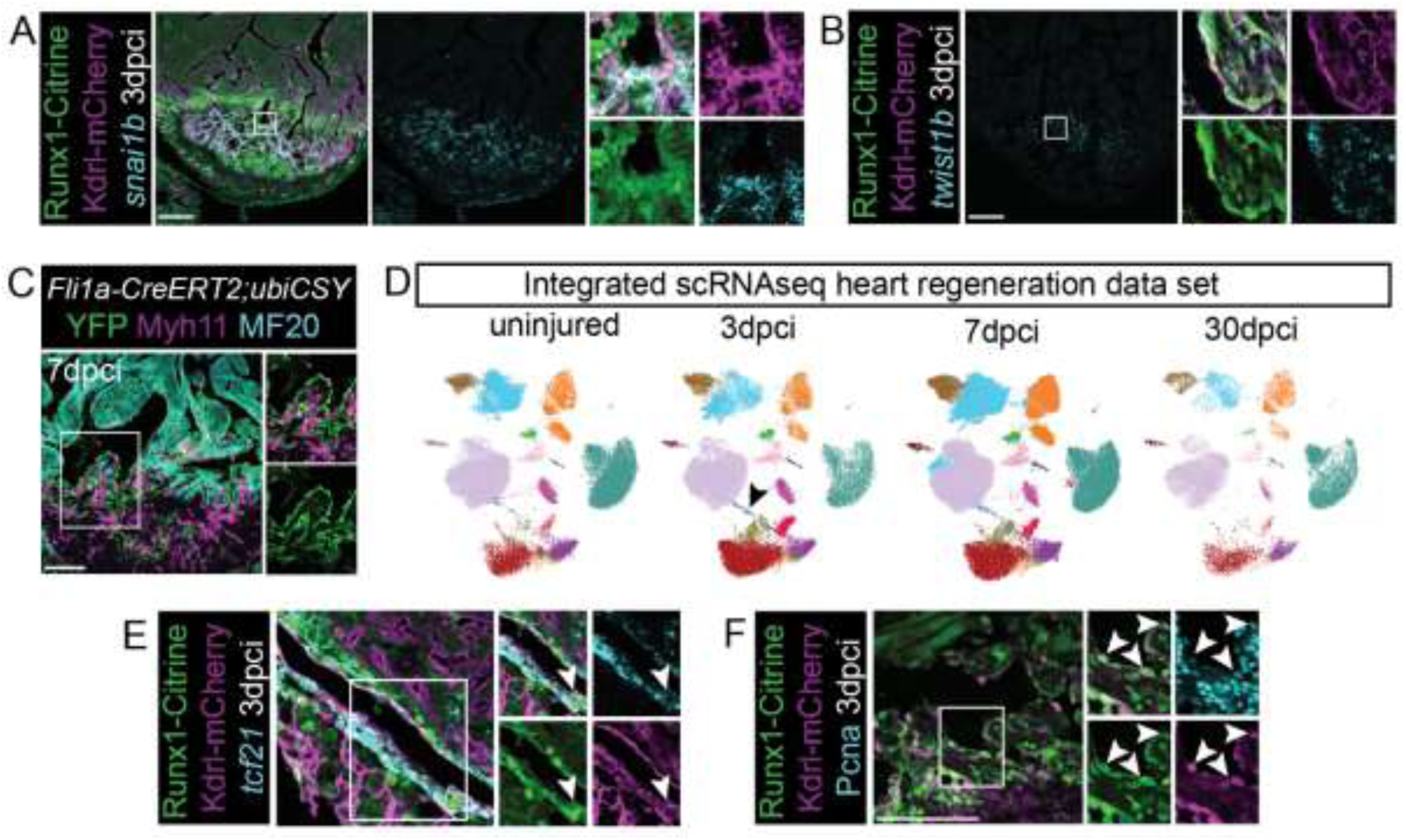
The injured endocardium is highly heterogeneous after injury. A-B, Immunohistochemistry showing labelling of Runx1:Citrine, Kdrl-mCherry and RNAscope staining for *snai1b* (A) or *twist1b* (B). C, *Tg(fli1a-creERT2;Tg(ubiCSY)* lineage tracing, immunohistochemistry for YFP (lineage traced endocardial cells), Myh11 and MF20 at 7dpci. The limited overlap of YFP and myh11 indicates the lack of endoMT into fully differentiated myofibroblasts. D, UMAP of the integrated zebrafish heart regeneration scRNAseq data set derived from the integration of 6 independent scRNAseq data sets at different timepoints after injury^9,30–34^ E, Immunohistochemistry showing labelling of Runx1:Citrine, Kdrl-mCherry combined with RNAscope staining for *tcf21*. F, Co-labelling of Runx1:Citrine with PCNA showcases the proliferating state of the Runx1:Citrine positive endocardium.

**Supplementary Figure 2.**
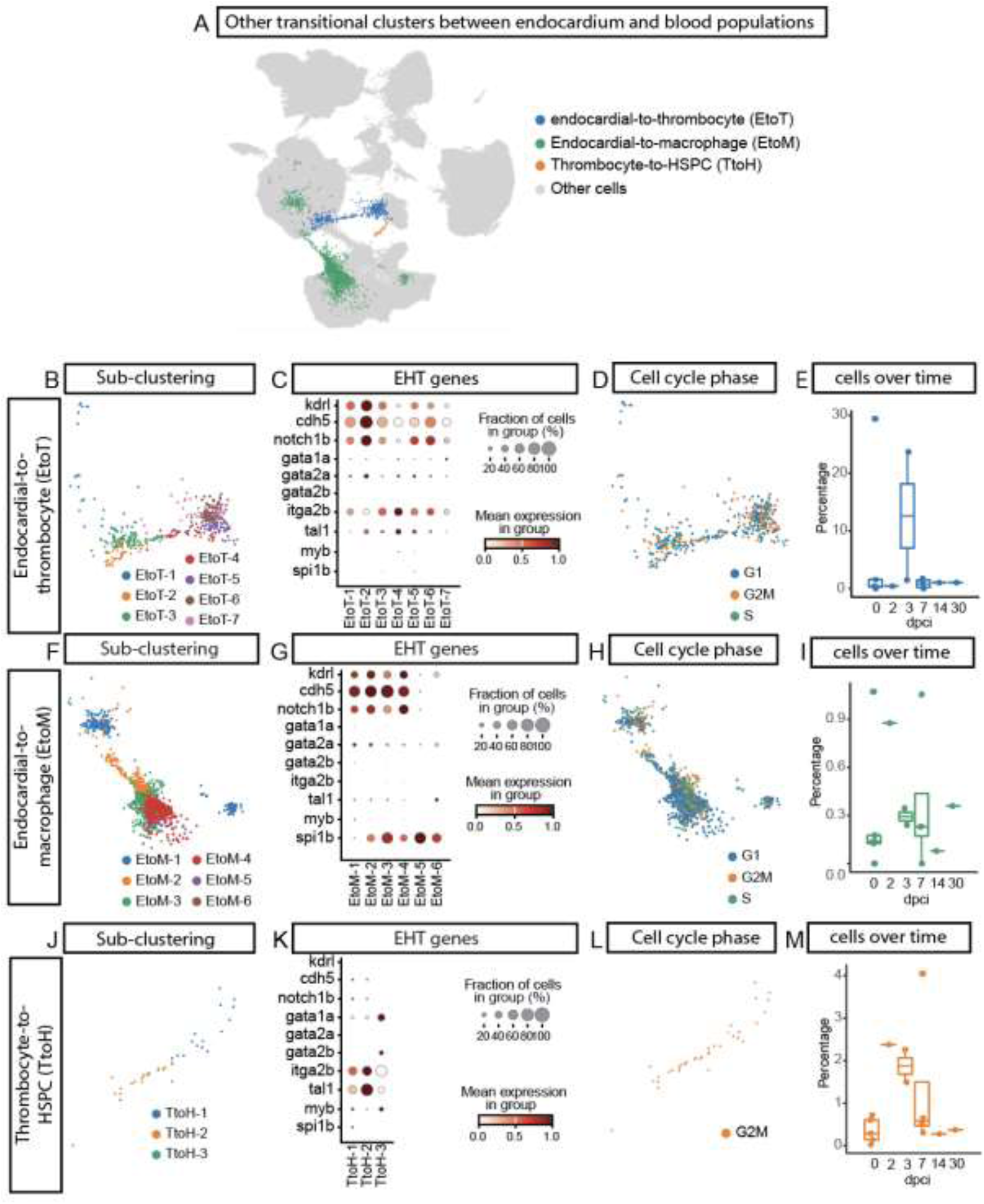
Analysis of possible additional transitional clusters between the endocardium and blood populations. A, UMAP of all cells of the integrated data set with additional cluster of interest highlighted. B, Sub-clustering of the bridging cluster connecting the endocardium directly with the thrombocyte cluster. C, Dotplot for endocardial and EHT gene expression in the EtoT subclusters from B, showing reduction of endocardial gene expression from the endocardial cells towards thrombocytes, but no increase in EHT gene expression towards the thrombocyte cluster, D, Cell cycle phase annotation for EtoT cluster. There was also no clear activation of the cell cycle. E, Cell number quantification in uninjured up to 30dpci in EtoT cluster. The EtoT bridge is injury-responsive. F, Subclustering of bridging cluster connecting the endocardium directly with the macrophage (EtoM) cluster. G, Endocardial and EHT gene expression analysis of the EtoM subclusters. Lack of gene expression suggesting EHT transition. H, Analysis of cell cycle activation in EtoM clusters, no clear activation of the cell cycle. I, Analysis of cell numbers in EtoM subclusters in uninjured and injured hearts over time. J, Subclustering of the very small bridging cluster connecting the thrombocyte and HSPC-like (TtoH) cluster. K, Gene expression analysis of endocardial to haematopoietic transition genes in the TtoH cluster. This bridge lacks endocardial gene expression, but expresses EHT genes such as *myb, gata2b* as well as *itga2b*. L, Cell cycle phase annotation for cells present in the of TtoH subcluster. Most cells in this bridging cluster were in the G2M phase of the cell cycle. M, Cell numbers over time of cells populating the TtoH cluster. The TtoH bridge is injury responsive.

**Supplementary Figure 3.**
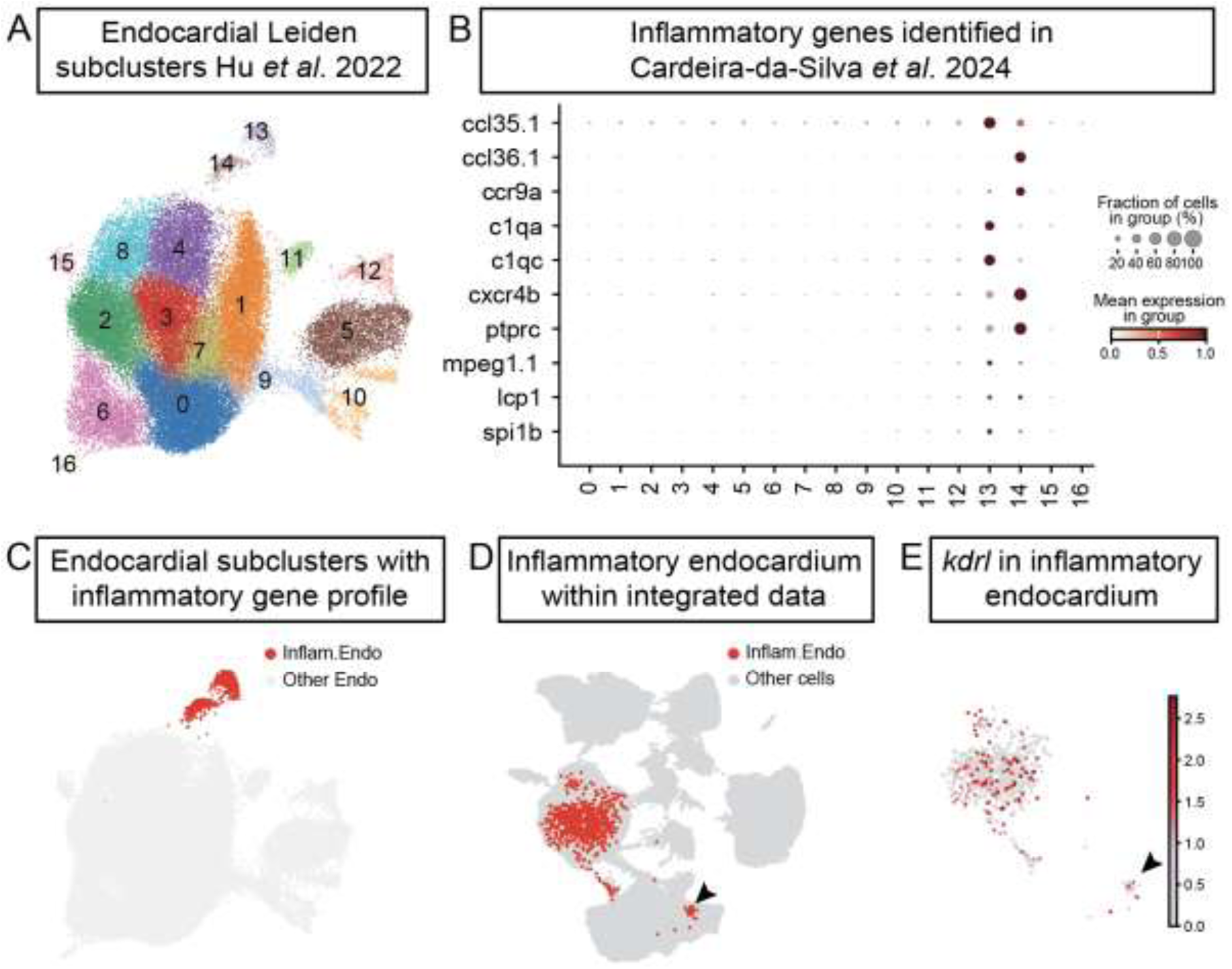
Sub clustering analysis of endocardial cells for inflammatory response genes. A, UMAP showing subclusters of uninjured and injured wild-type endocardial cells from Hu et al dataset ^34^, identifying 16 different subclusters. B, Dotplot for the expression of inflammatory genes as identified in Cardeira-da-Silva^45^ across the subclusters from A. C, UMAP highlighting the 2 endocardial subclusters with inflammatory gene expression. D, UMAP displaying the position of the inflammatory endocardial cells within the integrated data. Arrowhead points to cells with the inflammatory endocardial gene expression located within the T cell cluster of the integrated data set. E, *kdrl* expression within the inflammatory endocardial cells as present within the integrated data. The small population of cells within the T cell cluster expresses low levels of *kdrl* (arrowhead).

**Supplementary Figure 4.**
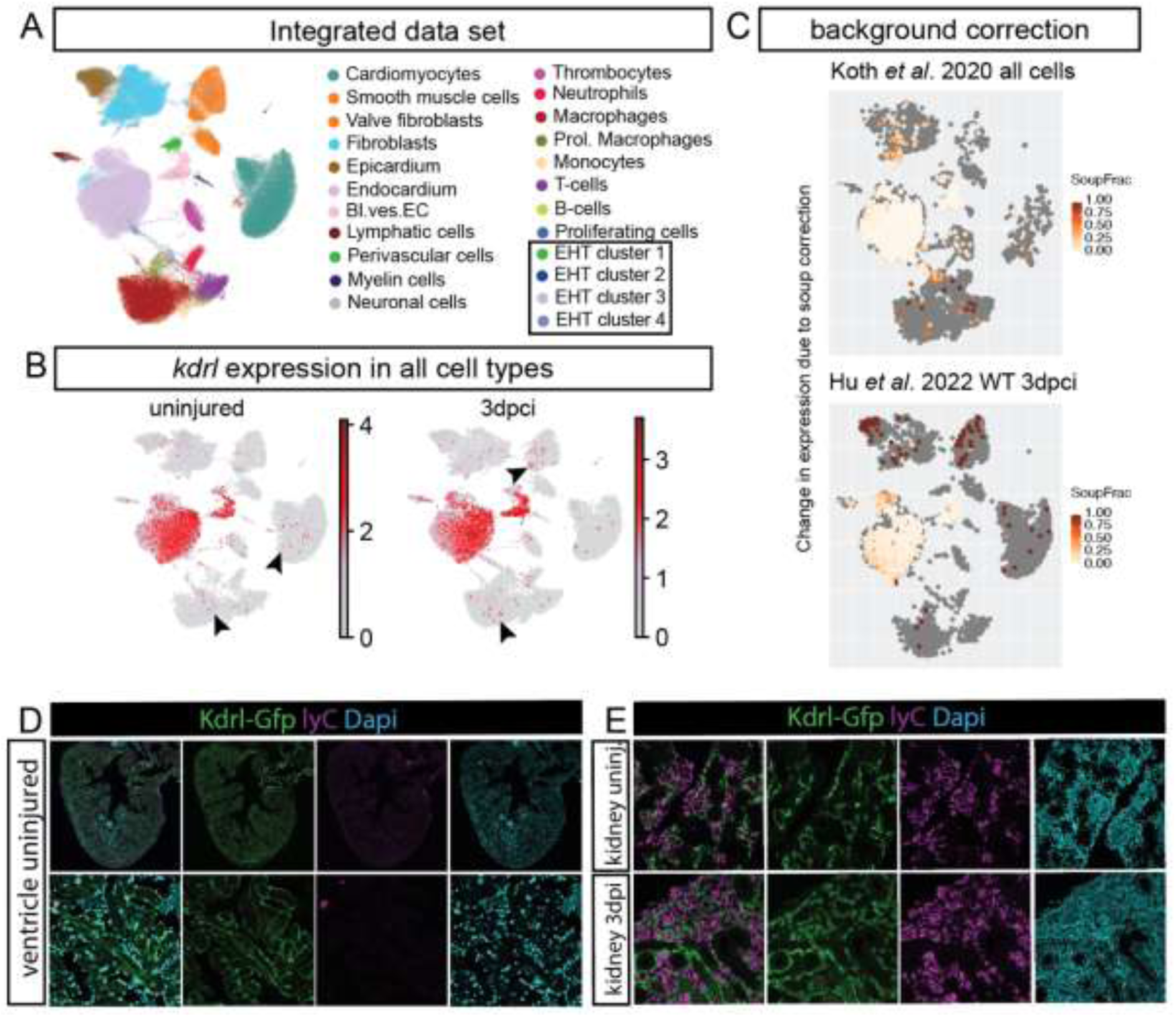
No evidence of Kdrl expression in leukocyte populations. A, UMAP of cell types present in the integrated data set. B, UMAP showing *kdrl* expression before injury and at 3dpci within the integrated data set. Arrowheads point to scattered *kdrl* expression in multiple cell type clusters C, UMAP showing the change in expression of *kdrl* due to soup correction to identify cells that can be considered background. This shows that almost all *kdrl* expression in the leukocyte populations can be considered as background levels. D, Immunohistochemical staining of uninjured ventricles of Kdrl-Gfp hearts for lyC positive blood cells showing absence of Kdrl-positive circulating cells. E, Immunohistochemistry for Kdrl-GFP and LyC in the head kidney before and at 3dpci of the heart showing lack of overlap between the two markers.

**Supplementary Figure 5.**
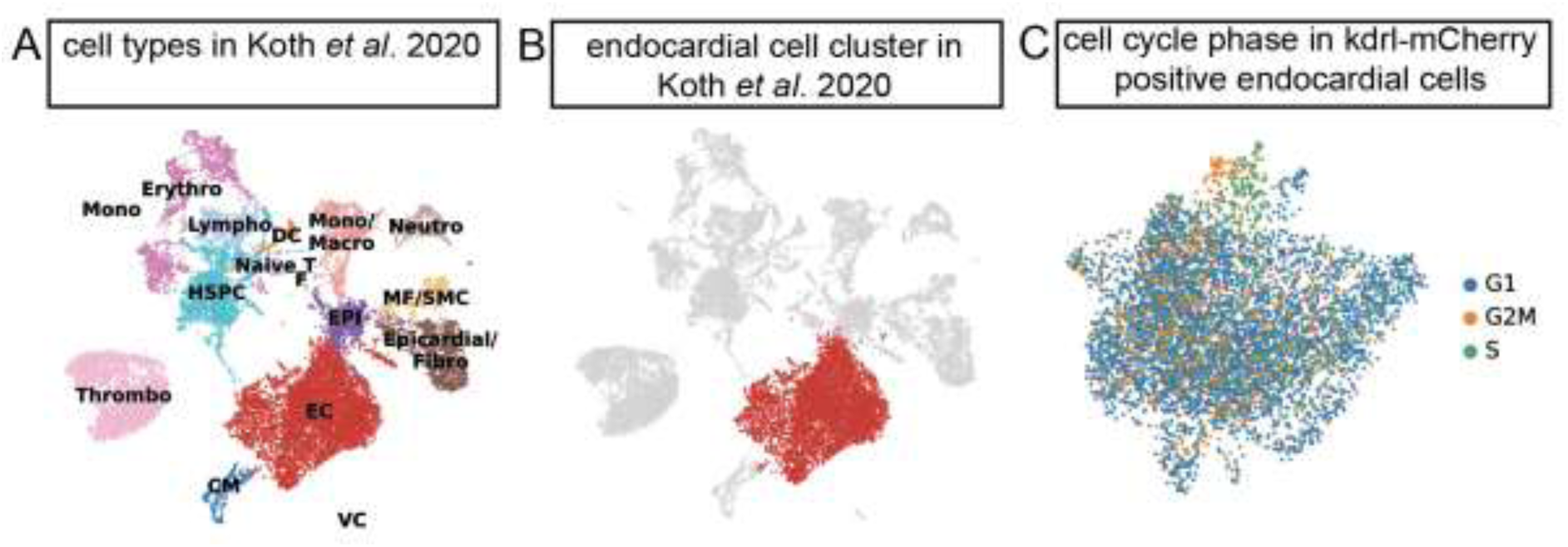
Subclustering analysis of Koth et al populations. A. UMAP plot of all cells analysed in Koth et al. 2020^9^ re-coloured by cell type annotation. B. UMAP of the global cell population displaying the endocardial cluster of interest which have been isolated for subclustering. C. UMAP plot of the isolated double positive kdrl-mCherry endocardial cell cluster from B annotated for cell cycle phase.

**Supplementary Figure 6.**
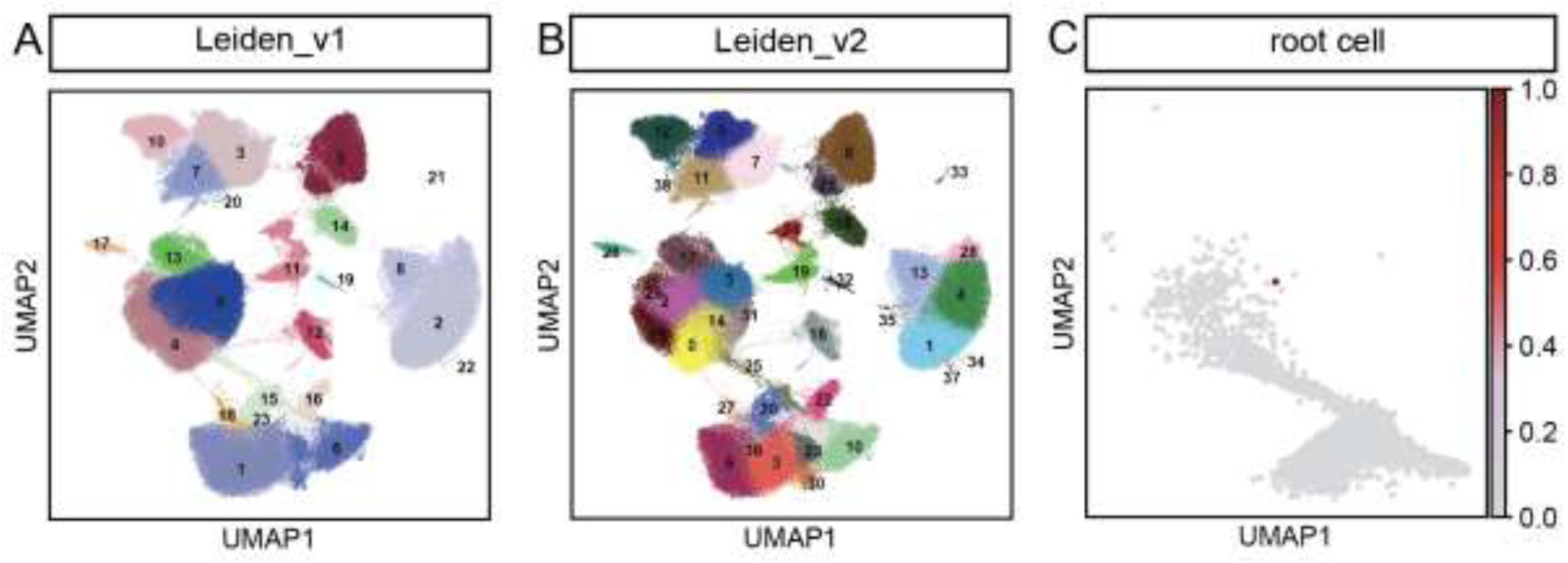
Details for scRNA-seq analysis. A-B, UMAP showing Leiden clustering of all cells with the resolution set to 0.8 (Leiden_v1) or 2.0 (Leiden_v2). C, UMAP showing the root cell selected for pseudotime analysis as shown in Figure 4E-F.

